# A Human Neuron Alzheimer’s Disease Model Reveals Barriers to Senolytic Translatability

**DOI:** 10.1101/2025.01.28.635165

**Authors:** Chaska C. Walton, Ellen Wang, Suckwon Lee, Cynthia J. Siebrand, Nicholas J. Bergo, Julie K. Andersen

## Abstract

Therapeutic successes in mouse models of Alzheimer’s disease (AD) largely fail to translate into clinical trials, with experimental drugs rarely validated in human models before being administered to humans. To address this, we developed an accessible method for long-term culture of commercially available primary human neurons and astrocytes, along with an amyloid-beta 1-42 (Aβ)-based *in vitro* AD model. Using this system, we evaluated two senolytic regimens previously shown to be effective in AD mouse models—Navitoclax and Dasatinib plus Quercetin (DQ)—and the natural killer cell line NK92 for emerging immune-mediated senescent cell ablation therapies. NK92 cells preferentially—but not exclusively—targeted Aβ-treated neurons and astrocytes with senescent-like phenotypes. DQ demonstrated a safe profile for human neurons, but Navitoclax exhibited non-selective neurotoxicity. These findings highlight risks of Navitoclax and NK-based interventions and underscore the critical need for human-relevant models in the AD drug-development pipeline to improve safety and clinical translatability.

## Introduction

The Food and Drug Administration (FDA) has approved three Aβ-clearing antibodies that, while effective at reducing biological markers of pathology, are yet to robustly stop clinicopathological AD progression^1–4^. Historically, a paradoxical disconnect has existed between positive results in preclinical research and the extraordinarily high failure rate in AD clinical trials^5–7^. This highlights two critical bottlenecks: the need for new therapies and the lack of translation from research in animal models to humans.

In this context, cellular senescence has emerged as a novel druggable target in AD^8^. Senescent cells, which accumulate with age, adopt a harmful senescence-associated secretory phenotype (SASP) that drives inflammation and contributes to tissue damage^9,10^. The selective elimination of senescent cells has demonstrated beneficial effects in preclinical studies, initially in mouse models of Parkinson’s disease^11^. Subsequently senolytics, a class of drugs used to eliminate senescent cells^12^, have shown promise in mitigating AD pathology in rodent models^13–17^ and have progressed to clinical trials for AD^8^.

Senolytics are broadly categorized under a unifying umbrella but demonstrate diverse mechanisms of action^18^. Perhaps unsurprisingly, the use of different senolytics in different models of AD-like pathology has resulted in conflicting evidence regarding which central nervous system (CNS) cell types—neurons, astrocytes, microglia, or oligodendrocyte progenitor cells (OPCs)—undergo deleterious cellular senescence which can be targeted by senolytics^13–18^. Indeed, different senolytics have been reported to eliminate distinct, non-overlapping senescent non-CNS cell populations^18^.

Laboratory-generated human neurons, such as those derived from induced pluripotent stem cells (iPSCs), are powerful tools commonly used to address species-specific differences inherent in the use of rodent models^19,20^. However, the developmental origins and cellular characteristics of laboratory-generated neurons differ fundamentally from those of bona fide human CNS neurons^21^. As mentioned above, senolytics exhibit cell-type-specific toxicity^18^. As a result, the ability of senolytic profiles observed in such laboratory-generated neurons to translate effectively to human neurons remains uncertain.

Human primary neuron cultures, derived directly from human CNS tissue, offer a biologically relevant model for studying age-related diseases including AD^21^. However, long-term culture protocols necessary to model the adult brain require access to specialized medical facilities^22–25^. This reliance has likely limited their widespread use, confining these approaches to a small number of laboratories over several decades^26–33^. Consequently, the toxicity and mechanisms of action of AD drugs, including senolytics, on human neurons often remain untested prior to use in clinical trials.

Here, we demonstrate that long-term culturing of mixed primary human neuron and astrocyte cells derived from commercially available cryopreserved preparations can be easily maintained over 100 days *in vitro* (DIV) and include neurons expressing key neuronal markers. Using this system, we evaluated the two senolytic drug regimens previously preclinically explored as AD interventions—Navitoclax (NAV) and the senolytic cocktail DQ—in an *in vitro* model of Aβ-induced AD pathology^8^. To benchmark NAV and DQ, we compare these senolytics to a model of physiological senescent cell ablation using the natural killer (NK) cell line NK92, modeling immune system-mediated clearance^34,35^. The results for NK92 co-culture experiments are consistent with the preferential ablation of senescent-like populations amidst non-selective elimination of healthy-control cells. While DQ did not show neurotoxicity, it moderately impacted astrocyte viability in a non-selective manner. In contrast, NAV exhibited dose-dependent toxicity in both healthy and Aβ-treated neurons. These findings underscore the need to refine senolytic strategies in human-relevant models of AD.

## Results

### Characterization of long-term human neuronal-astrocytic cultures

Cells extend processes (Videos 1, 2), undergo nuclear migration (Video 3), and cell division (Videos 3, 4) within 48 hours after plating. Based on morphology, different cell types are identifiable shortly after plating (Video 5). Neurons positive for the neuronal markers NeuN and MAP2 as well as GFAP-positive astrocytes are present early on (Fig. 1a). These cultures can be easily maintained for 100 days *in vitro* (DIV) (Fig. 1a, b) and we have successfully cultured up to 200 DIV (Sup. Fig. 1a); the upper age limit for these cultures is currently unknown. Human primary neurons present characteristic markers including a dense mesh of tau 3R-positive axons (Fig. 1c), positive immunostaining for the pan-tau marker tau-5 (Sup. Fig. 1b) and for Ankyrin-G (Sup. Fig. 1a, c), a marker for axon initial segments^36^. We observed several unexpected phenotypes in these cultures, with arguably the most striking being their extended viability (Sup. Fig. 1a). Another unexpected finding was that NeuN staining was not restricted to perinuclear regions but also extended into processes during maturation (Fig. 1a, b). We also detected soma staining with MN423 (Fig. 1c), a tau antibody purportedly exclusively recognizing the pronase-resistant core of tau paired helical filaments (PHF), but not full-length human tau^37,38^. Additionally, some axons were positive for MC1 antibody (Sup. Fig. 1d), which was previously thought to exclusively detect pathological tau conformations associated with neurofibrillary tangles^39^.

**Figure 1.**
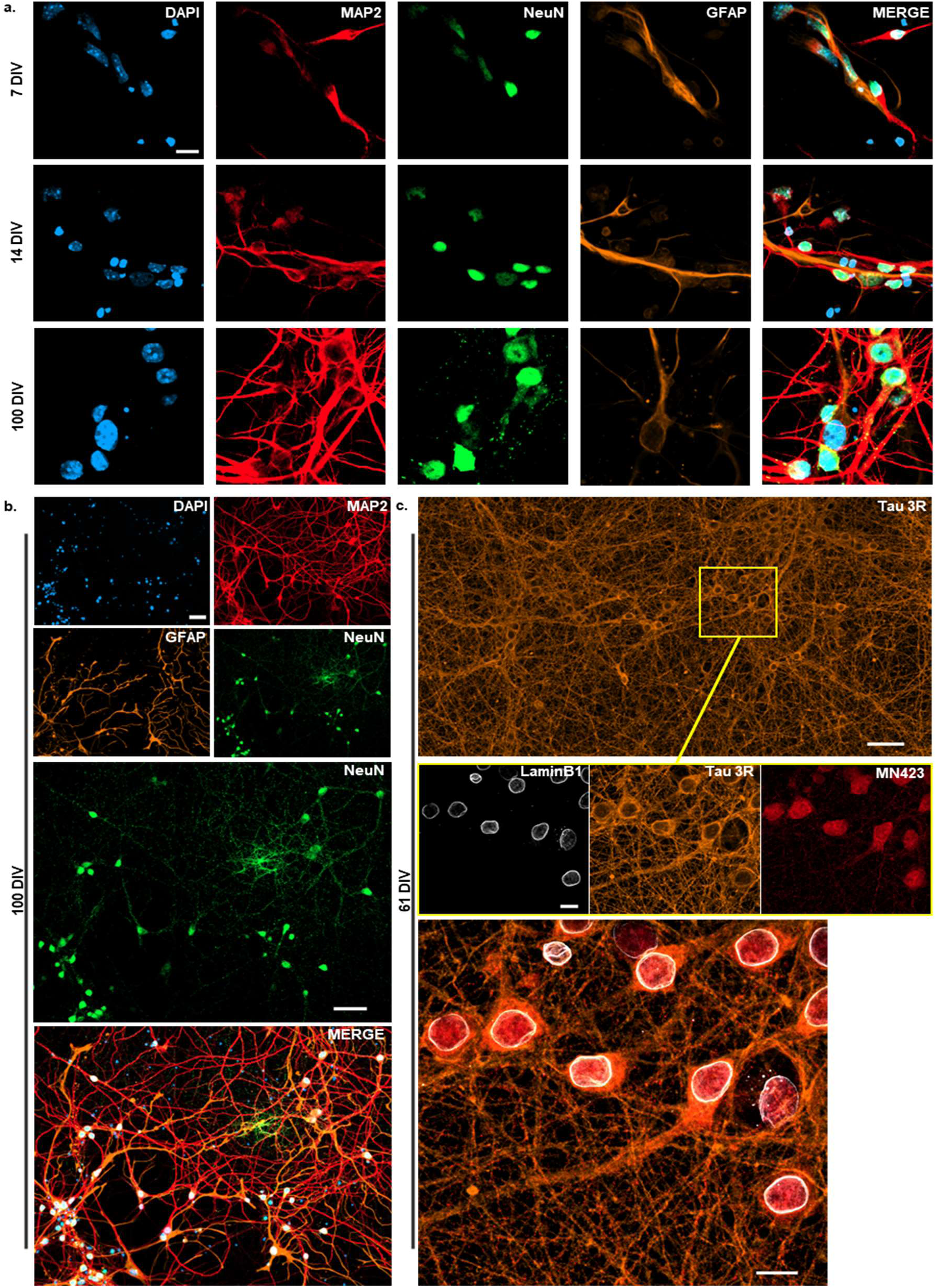
Development of neuronal and astrocytic markers in human primary neurons during maturation. (**a**) Super-resolution confocal images of cultures at 7, 14, and 100 DIV. DAPI (nuclei), MAP2 (dendrites), NeuN (neuron somas), and GFAP (astrocytes). The increasing complexity of MAP2 dendritic arbors reflects neuronal maturation. At 100 DIV, NeuN puncta extends beyond somatodendritic regions. 63x objective with 1.7x zoom. Scale bar = 10 µm. (**b**) Confocal tile scan of 100 DIV cultures with immunostaining as in (**a**). 20x objective with 1x zoom. Scale bar = 50 µm. (**c**) Confocal tile scan (top) of 61 DIV cultures with tau stained by tau 3R antibody. The detailed images (middle) focus on a region of the tile scan (yellow box) with immunostaining for LaminB1 (nuclear lamina), Tau 3R (axons), and MN423 (tau PHF). The enlarged image (bottom) merges LaminB1, tau 3R, and MN423. 40x objective with 2x zoom. Scale bars: tile scan (top) = 50 µm; detail images (middle) and merged images (bottom) = 10 µm.

Dendritic processes demonstrated a marked increase in number and complexity over time, as evidenced by MAP2 immunostaining (Fig. 1a, b). To further investigate neuronal maturation and assess synaptic development, we performed immunocytochemistry (ICC) for the presynaptic marker synapsin I (synapsin) and postsynaptic marker Homer1 (Fig. 2a), as well as synapsin and the postsynaptic marker PSD95 (Sup. Fig. 2) at 14, 28, and 43 DIV. Synapsin is a vesicle-associated protein localized at axon terminals, while Homer1 and PSD95 are scaffolding proteins found at the postsynaptic density^40,41^.

**Figure 2.**
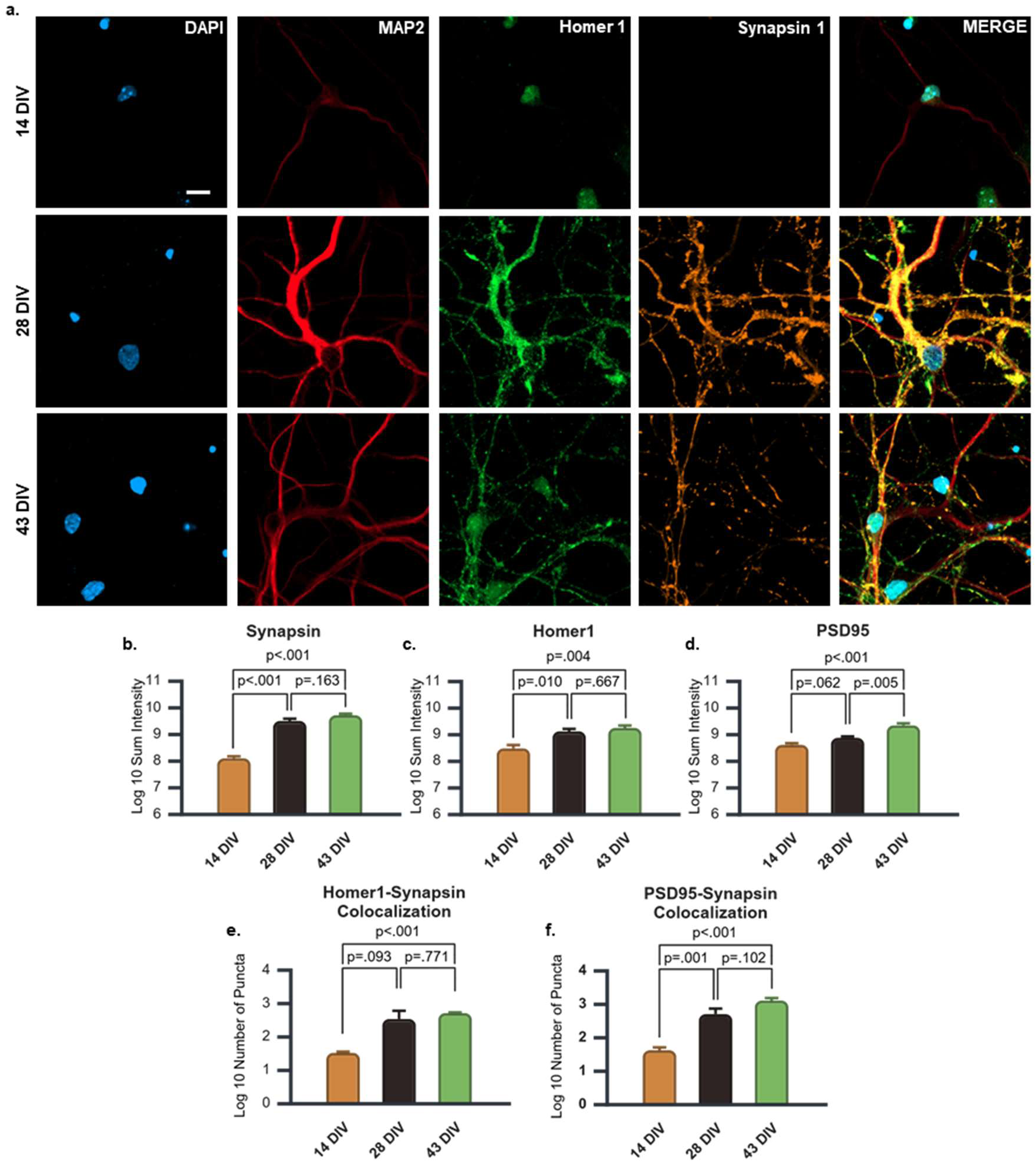
Synaptic markers in human primary neurons during maturation. (**a**) Representative super-resolution confocal images from cultures at 14, 28, and 43 DIV with staining for DAPI, MAP2, Homer1 (postsynaptic marker), and Synapsin (presynaptic marker). 63x objective with 2x zoom. Scale bar = 10 µm. (**b**) Quantification of log10-transformed total Synapsin puncta signal intensity at 14, 28, and 43 DIV. Three biological replicates, each with 2 technical replicates. (**c, d**) Quantification of log10-transformed total (**c**) Homer1 and (**d**) PSD95 puncta signal intensity at 14, 28, and 43 DIV. (**e, f**) Quantification of log10-transformed number of colocalized puncta of (**e**) Homer1-Synapsin and (**f**) PSD95-Synapsin at 14, 28, and 43 DIV. (**c-f**) Three biological replicates. (**b-f**) Significance, *p* < 0.05. Graphs display mean ± s.e.m.

A one-way ANOVA revealed significant changes in synapsin expression with maturation (*p* < 0.001; see Table 1a for omnibus test results). Multiple comparisons showed that synapsin intensity at 14 DIV was significantly lower than at 28 (*p* < 0.001) and 43 DIV (*p* < 0.001; Fig. 2b) However, the difference between 28 DIV and 43 DIV was not statistically significant (*p* < 0.163; Fig. 2b). Likewise, significant differences were also observed for Homer1 expression across DIV (*p* = 0.004; Table 1b), with significant increases in intensity between 14 and 28 DIV (*p* = 0.010) and between 14 and 43 DIV (*p* = 0.004), but not between 28 and 43 DIV (*p* = 0.667; Fig. 2c). These findings indicate a significant increase in both pre and postsynaptic markers between 14 and 28 DIV and between 14 and 43 DIV, reflecting robust synaptic development during early neuronal maturation, with a plateau in development observed between 28 and 43 DIV.

As per the analysis of PSD95 with maturation, there were also highly significant differences between DIV of culture (*p* < 0.001; Table 1c). A non-significant trend for increased PSD95 was observed between 14 and 28 DIV (*p* = 0.062), which was significant when comparing 14 and 43 DIV (*p* < 0.001) as well as 28 and 43 DIV (*p* = 0.005; Fig. 2d). These findings suggest a progressive increase in PSD95 expression, indicative of ongoing postsynaptic development between 14 and 43 DIV.

To further characterize synaptic maturity, we assessed the colocalization of Homer1 with Synapsin and PSD95 with Synapsin for each time point. Colocalized Homer1 and Synapsin puncta showed significant differences (*p* < 0.001; Table 1d), with a non-significant trend towards increased colocalization from 14 to 28 DIV (*p* = 0.093) and a significant increase between 14 and 43 DIV (*p* < 0.001; Fig. 2e). The differences between 28 and 43 DIV were not significant (*p* = 0.771; Fig. 2e). As per colocalized PSD95 and Synapsin puncta, there were also highly statistically significant differences with maturation (*p* < 0.001; Table 1e). There was a significant increase from 14 to 28 DIV (*p* = 0.001) and from 14 to 43 (*p* < 0.001) DIV but not from 28 to 43 DIV (*p* = 0.102; Fig. 2f). These results indicate significant increases in Homer1-Synapsin and PSD95-Synapsin colocalization during early neuronal maturation, with a plateau observed between 28 and 43 DIV.

### NK92 co-culture results in cell killing compatible with the preferential elimination of senescent-like neurons and astrocytes in an Aβ *in vitro* model of AD

The immune system naturally eliminates senescent cells through senescence immune surveillance, primarily mediated by NK cells, CD8+ cytotoxic T-cells (CTT), and macrophages^34^. *In vitro* studies have utilized NK92 cells, a human NK cell line, to model the immune-mediated clearance of senescent cells^35^. To investigate the potential of senescence immune surveillance as a therapeutic for AD as well as to benchmark NAV and DQ, we co-cultured NK92 cells with primary neurons and astrocytes at 42 DIV for 24 hours and assessed whether there were differences in selective killing between Aβ(1-42)-treated (0.5 μM, 28 DIV) and vehicle-treated cells (Figure 3a).

**Figure 3.**
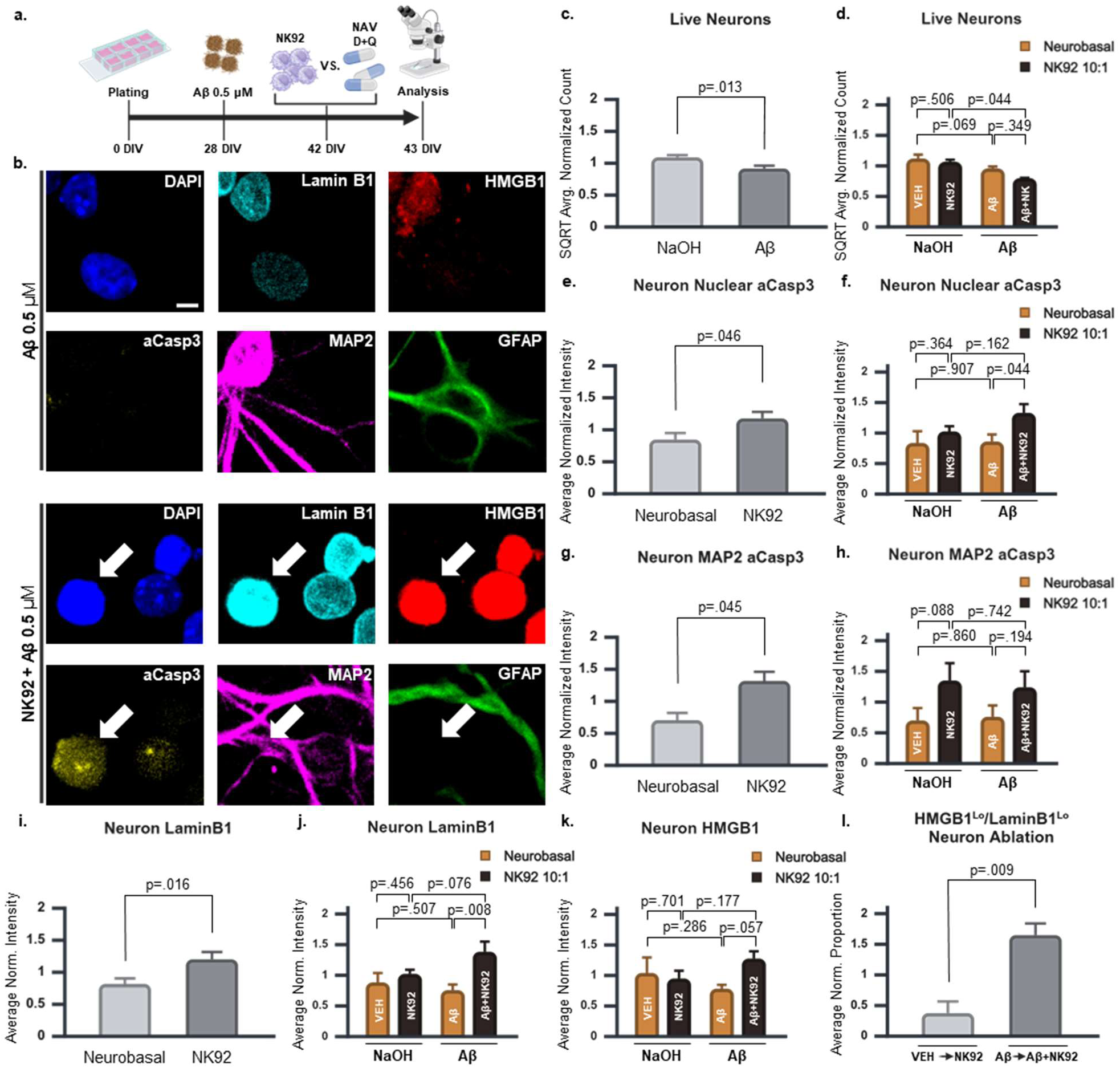
I*n vitro* AD model of senescence immune surveillance in human neurons. (**a**) Diagram depicting human *in vitro* model of AD to assess senescent cell ablation by NK92, NAV, and DQ. Primary mixed human neuron and astrocyte cultures were treated with either vehicle (NaOH) or 0.5 μM of Aβ preparations enriched in oligomers at 28 DIV. At 42 DIV, NK92 cells, NAV or DQ were added to the cultures. Cells were fixed for ICC at 43 DIV. (**b**) Representative linearly unmixed confocal spectral scans of Aβ (top 2 rows) and Aβ + NK92 (bottom two rows) groups (also see Sup. Fig. 3b). The white arrow identifies a putative NK92 cell that is MAP2^-^/GFAP^-^/aCasp3^+^. 40x objective with 1x zoom. Scale bar = 5 µm. (**c**) Aβ main effects and (**d**) Aβ x NK92 simple main effects on square root (SQRT) transformed average normalized viable neuron count. The data was SQRT-transformed to meet the assumption of normality (Table 2a). (**e**) NK92 main effect and (**f**) Aβ x NK92 simple main effects on nuclear aCasp3. (**g**) NK92 main effect and (**h**) Aβ x NK92 simple main effects on aCasp3 within MAP2 processes. (**i**) NK92 main effect and (**j**) Aβ x NK92 simple main effects on LaminB1. (**k**) Aβ x NK92 simple main effects on HMGB1. (**l**) Comparison of HMGB1^Lo^/LaminB1^Lo^ neuronal loss between VEH→NK92 and Aβ→Aβ+NK92, quantified as described in Sup. Fig 3e. (**c, e, g, i**) Significance, *p* < 0.05. (**d, f, h, j, k**). Significance, *p* < 0.025. (**l**) Significance, *p* < 0.0167. 3x Replicates. A total of 6111 neurons were analyzed. Graphs represent mean ± s.e.m.

First, we performed exploratory time-lapse imaging experiments to track live cell behavior in the absence of NK92 cells (Video 6) and in co-culture with NK92 cells for both non-Aβ (Video 7) and Aβ-treated cultures (Video 8; Sup. Fig. 3a). NK92 cells were viable in neurobasal medium, formed uropods, and actively interacted with target neurons and astrocytes irrespective of Aβ treatments (Videos 7 and 8). Having established the feasibility of NK92 co-cultures, we used confocal spectral scanning with linear unmixing to determine whether the effects of NK92 co-culture were consistent with selective senescent cell ablation. This imaging technique allowed us to simultaneously resolve the 6 channels resulting from immunocytochemistry (ICC) for DAPI to identify nuclei, HMGB1 and LaminB1 as markers of cellular senescence^10,42–44^, active caspase-3 (aCasp3) to identify apoptotic cells^45^, as well as MAP2 and GFAP to distinguish neurons and astrocytes, respectively (Fig. 3b; Sup. Fig. 3b, c; Video 9). Confocal tile scans covering several hundred square microns were acquired across three replicates for each treatment condition (Sup. Fig. 3d; Video 9).

For the statistical analyses of markers, two-way ANOVAs were run when the appropriate parametric assumptions were met. The criteria to establish significance and non-significant trends are described in the Materials and Methods section. Interactions and main effects are reported and graphed only if significant. Simple main effects with exact *p* values are always graphed. All statistical results can be found in the tables referenced in each analysis (see Materials and Methods for additional details). We first examined whether NK92 co-culture affected neuronal viability by analyzing the number of cells categorized as live neurons in each condition. A two-way ANOVA showed there was a significant main effect for Aβ treatment, revealing a reduction in the number of viable neurons with Aβ treatment (*p* = 0.013, Fig. 3c; Table 2a). The main effects (overall effects) of Aβ treatment assess the differences between the average viability of the VEH and NK92 group means and the average of the Aβ and Aβ+NK92 group means, reflected in Fig. 3c. Subsequent analysis of the simple main effects showed that when comparing Aβ+NK92 to NK92 groups the difference approached statistical significance (*p* = 0.044, Bonferroni-adjusted *p* < 0.025 for two simple main effects), with a large effect size (partial η² = 0.417), suggesting increased vulnerability of Aβ-treated neurons to NK92 cells (Fig. 3d; Table 2a). The simple main effects (condition-specific) analyses entail the specific comparisons between VEH, NK92, Aβ, and Aβ+NK92 treatments depicted in Fig. 3d.

In neurons, we separately assessed cytoplasmic and nuclear aCasp3 to distinguish initial activation from subsequent nuclear translocation during apoptosis^46,47^. Confocal images of aCasp3 showed that NK92 cells likely underwent apoptosis (Fig. 3b; Sup. Fig. 3b, c), so the analysis was restricted to live neurons and astrocytes positive for MAP2 and GFAP, respectively. For neurons, the main effects for NK92 were statistically significant, revealing increased nuclear aCasp3 with NK92 co-culture irrespective of Aβ treatment (*p* = 0.046, Fig. 3e; Table 2b). The main effects (overall effects) of NK92 treatments assess the differences between the average of the VEH and Aβ group means and the average of the NK92 and Aβ+NK92 group means, reflected in Fig. 3e. Condition-specific analysis suggested a non-significant trend toward higher nuclear aCasp3 in Aβ+NK92 cells compared to Aβ-only cells (*p* = 0.044), an effect absent when comparing NK92-only to vehicle-treated groups (*p* = 0.364, Fig. 3f; Table 2b). These results are consistent with preferential elimination of Aβ-treated neurons.

The analysis of aCasp3 within MAP2-positive neuronal processes also increased significantly with NK92 co-culture irrespective of Aβ treatment (*p* = 0.045, Fig. 3g; Table 2c), revealing non-selective effects. No significant differences or trends were found between treatment groups (Fig. 3h), indicating cytoplasmic aCasp3 was not preferentially accumulating in Aβ-treated cells. Notably, in confocal images, a clear aCasp3 signal was not visually discernible in MAP2-positive processes, suggesting that the apoptotic signal detected in the analysis reflects subtle changes. Overall, the analyses of nuclear (*p* = 0.046) and somatodendritic aCasp3 (*p* = 0.045) indicate a non-selective increase of cell ablation with NK92 co-culture, which may be more advanced in Aβ-treated neurons, as per a trend towards increased nuclear aCasp3 staining in Aβ+NK92 relative to Aβ (*p* = 0.044) that was not present in NK92 relative to VEH-control conditions (*p* = 0.364).

Next, we assessed whether there were changes in LaminB1 and nuclear HMGB1, which are reported to be reduced in a variety of cells undergoing cellular senescence^10,42–44^. NK92 co-culture significantly increased LaminB1 levels overall (*p* = 0.016, Fig. 3i; Table 2d), with a non-significant trend for the interaction with Aβ treatments (*p* = 0.090, Table 2d). For the two-way ANOVA, the interaction assessed whether the difference between VEH and NK92 were different from the difference between Aβ and Aβ+NK92. Said difference of differences is expected from selective or preferential senescent cell ablation. Gaining further insights into the interaction trend, follow-up analysis of condition-specific effects showed that NK92 co-culture significantly increased LaminB1 in Aβ-treated neurons (*p* = 0.008), but not in non-Aβ-treated neurons (*p* = 0.456, Fig. 3j; Table 2d). The effect size for Aβ-treated neurons (partial η² = 0.606) was much greater than that for vehicle-treated neurons (partial η² = 0.071), supporting a biologically relevant effect of NK92 cells on Aβ-treated neurons. Nuclear HMGB1 analysis did not reach significance, though the interaction term approached a non-significant trend (*p* = 0.101, Table 2e). Condition-specific comparisons between Aβ and Aβ+NK92 groups similarly showed an increase in population-wide nuclear HMGB1 that approached a non-significant trend (*p* = 0.057, Fig. 3k; Table 2e).

Our analysis for LaminB1 and HMGB1 could indicate underlying changes in neuron populations that simultaneously expressed low levels of LaminB1 and nuclear HMGB1 (HMGB1^Lo^/LaminB1^Lo^). To assess this, we identified HMGB1^Lo^/LaminB1^Lo^ neurons and calculated the loss of these populations from VEH to NK92 (VEH→NK92) and from Aβ to Aβ+NK92 (Aβ→Aβ+NK92), as described in Sup. Fig. 3e. A Bonferroni correction for the repeated use of HMGB1 and LaminB1 datapoints (3 tests) established the threshold for significance at *p* < 0.0167. The analysis revealed that the loss of HMGB1^Lo^/LaminB1^Lo^ neurons for Aβ→Aβ+NK92 was significantly higher than for VEH→NK92 (*p* = 0.009; Fig. 3l; Table 2f; Sup. Fig. 3e, f). This is consistent with the elimination of senescent-like cells identified by both low LaminB1 and low nuclear HMGB1.

We conducted a similar analysis for GFAP-positive astrocytes. The assessment of live astrocytes revealed a non-significant trend toward reduced viability in response to Aβ treatment (*p* = 0.098), an effect not observed with NK92 treatment (*p* = 0.141; Table 2g). Condition-specific analysis did not identify significant differences or trends (Fig. 4a; Table 2g), indicating lack of effects of NK92 on astrocyte live cell counts as analyzed. However, a two-way ANOVA revealed that NK92 co-culture significantly increased overall nuclear aCasp3 in astrocytes regardless of Aβ treatment (*p* = 0.009; Fig. 4b; Table 2h) and double GFAP/aCasp3-positive astrocytes were readily identifiable (Sup. Fig. 3c), indicating non-selective elimination was taking place to some degree.

**Figure 4.**
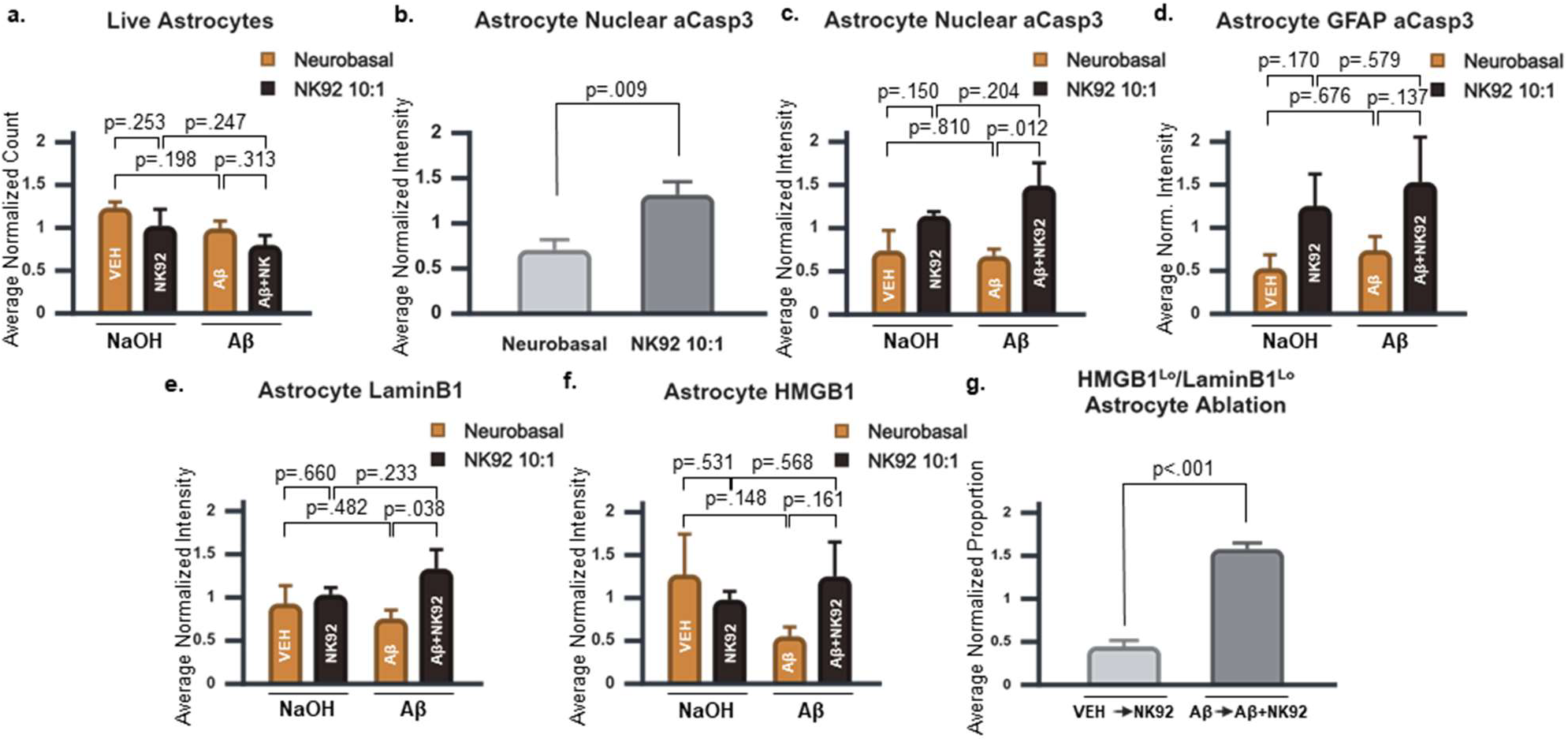
*In vitro* AD model of senescence immune surveillance in human astrocytes. (**a**) Aβ x NK92 simple main effects live astrocyte count. (**b**) NK92 main and (**c**) Aβ x NK92 simple main effects on nuclear aCasp3. (**d, e, f**) Aβ x NK92 simple main effects of (**d**) aCasp3 signal intensity within GFAP positive processes, (**e**) LaminB1 and (**f**) HMGB1. (**g**) Comparison of HMGB1^Lo^/LaminB1^Lo^ GFAP astrocytes between VEH◊NK92 and Aβ◊Aβ+NK92, as described in Sup Fig 3e. (**a, c, d, e, f**) Significance, *p* < 0.025. (**b**) Significance, *p* < 0.05. (**g**) Significance, *p* < 0.0167. 3x Replicates. A total of 1661 astrocytes were analyzed. Graphs represent mean ± s.e.m.

Reminiscent of the results for neurons (Fig. 3f), further condition-specific analysis of nuclear aCasp3 revealed a significant difference between Aβ and Aβ+NK92 treatment groups (*p* = 0.012) that was not observed between VEH and NK92 groups (*p* = 0.150; Fig. 4c; Table 2h), suggesting preferential elimination of Aβ-treated astrocytes was also taking place. The analysis of aCasp3 signal within GFAP-positive astrocytic processes showed NK92 co-culture only resulted in a trend towards higher cytoplasmic aCasp3 (*p* = 0.056; Table 2i). Although significant differences between conditions were not detected (Fig. 4d; Table 2i), the bar graph suggested that both NK92 and Aβ+NK92 conditions contributed to the non-significant trend observed for NK92 main effects.

The analysis of senescence biomarkers indicated a non-significant trend toward increased LaminB1 levels with NK92 co-culture (*p* = 0.071; Table 2j). Further condition-specific comparisons showed a non-significant trend of increased LaminB1 levels in Aβ+NK92 compared to Aβ alone (*p* = 0.038; Fig. 4e; Table 2j), possibly indicating that the trend for the main effects is mainly driven by Aβ+NK92. Nuclear HMGB1 analysis did not reveal significant effects or trends in main effects or interactions (Fig. 4f; Table 2k). However, when combining LaminB1 and HMGB1 data as performed previously for neurons (Sup. Fig. 3e), the analysis showed a highly significantly greater loss of HMGB1^Lo^/LaminB1^Lo^ astrocytes in Aβ→Aβ+NK92 compared to VEH→NK92 (*p* < 0.001; Fig. 4g; Table 2l; Sup. Fig. 4).

Our results indicate a biologically relevant preferential elimination of Aβ-treated neurons and astrocytes with senescent-like phenotypes, yet NK92 cells also exhibited cytotoxicity towards vehicle-control neurons and astrocytes. This effect was particularly pronounced in two co-cultures that were excluded from analysis due to severe cytotoxicity. Compared to their respective VEH (Video 10) and Aβ (Video 11) controls, these NK92 co-cultures resulted in the elimination of most cells regardless of whether the cells had been treated with the vehicle for Aβ (Video 12) or Aβ (Video 13). Thus, our results are aligned with senescence immune surveillance of Aβ-treated cells in the context of non-selective cytotoxicity.

### Navitoclax toxicity is not compatible with the preferential elimination of senescent cells in Aβ *in vitro* model of AD

NAV inhibits the Bcl-2 family of anti-apoptotic proteins and is extensively used as a senolytic compound^18,48–50^. It was originally reported to selectively kill mouse astrocytes and spare mouse neurons to achieve therapeutic effects in a transgenic mouse model of tauopathy^13^. To gain further insights into the effect of NAV on human neurons and astrocytes, cultures were treated with the vehicle for Aβ or Aβ 0.5 μM at 28 DIV as performed previously for NK92 cell experiments (Fig 3a). NAV was added at 14 days post-treatment (DPT) for 24 hours and cells were fixed for ICC. For our initial experiments we chose NAV treatments at 4 and 8 μM.

Regardless of whether cells were treated with Aβ or not, aCasp3 positive processes were evident (Sup. Fig. 5a). Although we did not detect double positive aCasp3 and MAP2 cells, we did observe aCasp3 positive cell remnants with long processes resembling dendrites and axons. To gain further insight into the morphology of neuron-like apoptotic remnants, we created a new aCasp3 channel (aCasp3*) that enhanced aCasp3 signal in cells that had already undergone cell death as a function of their loss for the rest of the markers, as described in Sup. Fig. 5b. This new channel revealed extensive aCasp3* positive processes that were consistent with axonal and dendritic morphology (Sup. Fig. 5b, c). In addition to aCasp3, neuronal loss could also be inferred from the visually evident deterioration of MAP2 positive processes in NAV-treated cultures irrespective of Aβ treatment (Sup. Fig. 5d). GFAP positive puncta, which could be remnants of astrocytic processes, also appeared after NAV treatment but the effects on astrocyte viability were not as evident (Sup. Fig. 5d).

We used the processed aCasp3* channel (Sup. Fig. 5b) to determine whether there were differences in apoptotic cell death between treatment conditions. To quantify apoptotic cell death, we calculated the ratio of live cells to aCasp3*-positive cell remnants negative for MAP2 and GFAP, as described in Sup. Fig. 5e. Of note, this analysis was not feasible for NK92 experiments, as there was no way to distinguish between the apoptotic cell remnants of neurons, astrocytes, and NK92 cells (Fig. 3b; Sup. Fig. 3b, c). The analysis showed that NAV treatments significantly increased apoptotic cell death regardless of Aβ treatment (*p* < 0.001; Sup. Fig 5f; Table 3a), reflecting lack of selectivity. We also assessed differences in the volume of MAP2 and GFAP-positive processes per unit area, revealing a significant reduction in neuronal processes at all concentrations (*p* < 0.001; Sup. Fig. 5g; Table 3b) and a significant reduction in astrocytic processes at 4 μM NAV (*p* = 0.038; Sup. Fig. 5h; Table 3c) relative to VEH that was independent of Aβ exposure, corroborating the lack of selectivity at these concentrations.

Given the toxicity observed at 4 and 8 μM NAV, we also assessed NAV at 0.5 μM, a concentration within the lower range of those tested *in vitro*^18^. The experimental design mirrored that employed in the NK92 and high-concentration NAV experiments, but with NAV at 0.5 µM (Fig. 3a). MAP2 and GFAP negative apoptotic cell remnants could be identified in the original aCasp3 channel under NAV treatment irrespective of the presence of Aβ (Fig. 5a). Following the same approach used in the 4 μM and 8 μM NAV treatment experiments (Sup. Fig. 5b), we generated an aCasp3* channel from the original aCasp3. As in higher-dose treatments, aCasp3* signal was also evident in NAV-treated conditions with or without Aβ (Sup. Fig. 5i). However, unlike for higher NAV concentrations, the effects of NAV at 0.5 μM on MAP2 and GFAP processes did not indicate obvious impacts on neuron and astrocyte populations (Sup. Fig. 5j). Together, this suggested a moderate effect of NAV at 0.5 μM.

**Figure 5.**
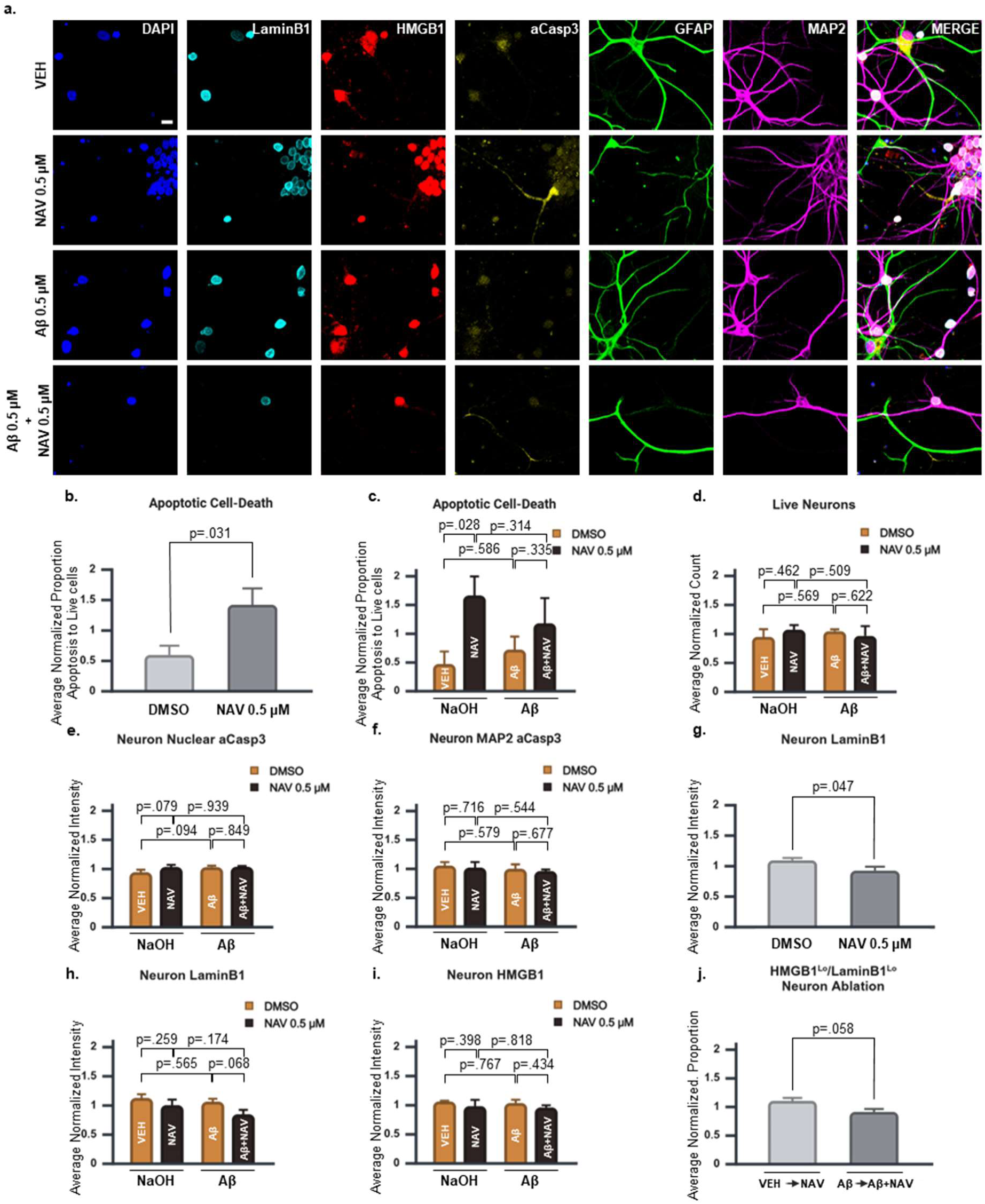
Assessment of NAV 0.5 μM on human primary neurons. (**a**) Representative linearly unmixed confocal spectral scans of all treatment conditions of NAV 0.5 µM experiments. 40x objective with 1x zoom. Scale bar = 10 µm. (**b**) NAV main effects and (**c**) Aβ x NAV simple main effects on apoptotic cell death, quantified as described in Sup. Fig. 5e. (**d, e, f**) Aβ x NAV simple main effects on (**d**) live neuron count, (**e**) nuclear aCasp3 and (**f**) aCasp3 within MAP2 processes. (**g**) NAV main effects and (**h**) Aβ x NAV simple main effects on LaminB1. (**i**) Aβ x NAV simple main effects on HMGB1. (**j**) Comparison of HMGB1^Lo^/LaminB1^Lo^ neuronal loss for VEH◊NAV and Aβ◊Aβ+NAV, as described in Sup Fig 3e. (**b, g**) Significance, *p* < 0.05. (**c, d, e, f, h, i**) Significance, *p* < 0.025. (**j**) Significance, *p* < 0.0167. 3x Replicates. A total of 9840 neurons were analyzed. Graphs represent mean ± s.e.m.

For apoptotic cell death, a two-way ANOVA revealed that NAV at 0.5 μM significantly increased apoptosis overall (*p* = 0.031; Fig. 5b; Table 4a). Unexpectedly, further condition-specific analysis suggested a non-significant trend toward increased apoptotic cell death in NAV-treated VEH cells (*p* = 0.028), which was not observed in Aβ-treated cells (*p* = 0.335; Fig. 5c; Table 4a). However, no significant differences were detected in the viable neuron population remaining after NAV treatment (Fig. 5d; Table 4b), nuclear aCasp3 levels (Fig. 5e; Table 4c), or aCasp3 within MAP2-positive processes (Fig. 5f; Table 4d). These findings suggest that the apoptotic cell remnant may not have been neurons or the effects of NAV 0.5 μM were mild enough to avoid detection by our metrics.

Surprisingly, NAV treatment significantly reduced the population mean of LaminB1 in neurons overall (*p* = 0.047; Fig. 5g; Table 4e), opposite to what would be expected from the elimination of a senescent-like neuron population characterized by low LaminB1 expression. Condition-specific analysis did not reveal additional trends (Fig. 5h; Table 4e), indicating reduction in LaminB1 was taking place irrespective of Aβ treatment. HMGB1 analysis did not yield significant effects or trends (Fig. 5i; Table 4f). Similarly, the assessment of HMGB1^Lo^/LaminB1^Lo^ neurons showed no significant differences between VEH→NAV and Aβ→Aβ+NAV (*p* = 0.058; Fig. 5j; Table 4g; Sup. Fig. 5k). Thus, any undetected preferential killing associated with NAV treatment would not involve the selective elimination of HMGB1^Lo^/LaminB1^Lo^ neurons.

In terms of the effects on astrocytes, there were no observable significant results. The analysis of live astrocytes (Fig. 6a; Table 4h), aCasp3 signal within astrocyte nuclei (Fig. 6b; Table 4i), and aCasp3 within GFAP positive processes (Fig. 6c; Table 4j) did not reveal significant effects. Similarly, the analysis of astrocytic LaminB1 (Fig. 6d; Table 4k), HMGB1 (Fig. 6e; Table 4l), and HMGB1^Lo^/LaminB1^Lo^ astrocytes (Fig. 6f; Sup. Fig. 6; Table 4m) did not provide statistical support for selective elimination of cells by NAV.

**Figure 6.**
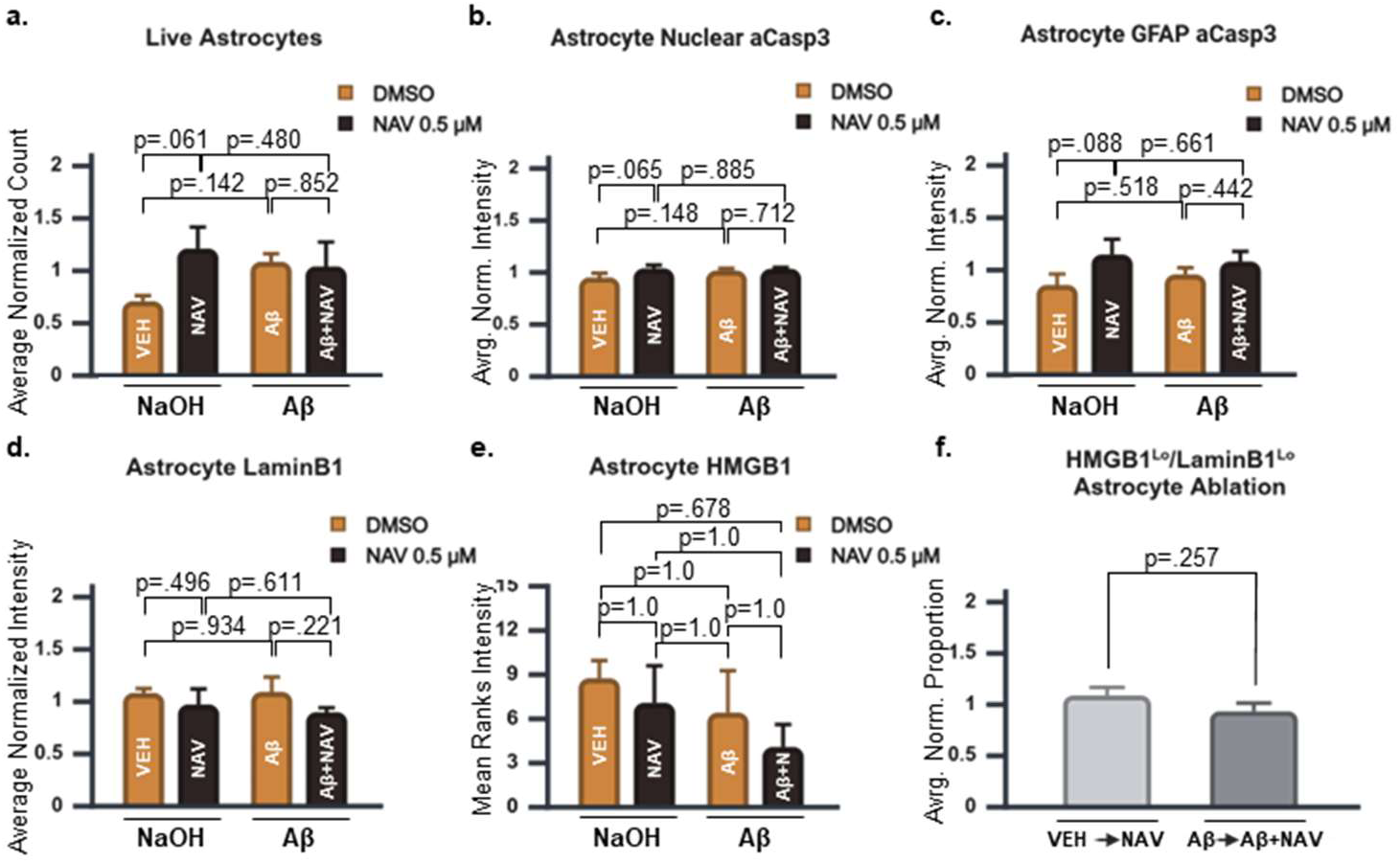
Assessment of NAV 0.5 μM on human primary astrocytes. (**a, b, c, d**) Aβ x NAV simple main effects on (**a**) live astrocyte count, (**b**) nuclear aCasp3, (**c**) aCasp3 within GFAP processes, and (**d**) LaminB1 signal. (**e**) Mean ranks for nuclear HMGB1. (**f**) Comparison of HMGB1^Lo^/LaminB1^Lo^ astrocyte loss for VEH→NAV and Aβ→Aβ+NAV, as described in Sup Fig 3e. (**a, b, c, d**) Significance, *p* < 0.025. (**e**) Significance, *p* < 0.05. (**f**) Significance, *p* < 0.0167. 3x Replicates. A total of 2216 astrocytes were analyzed. Graphs represent mean ± s.e.m.

Overall, NAV treatment at high concentrations (4 and 8 μM) resulted in non-selective elimination of both neurons and astrocytes. The analysis of apoptotic cell death following lower-concentration NAV treatment (0.5 μM) further supported the lack of selectivity. Cell type-specific viability and apoptosis metrics, however, did not clearly indicate whether neurons or astrocytes were preferentially affected at this lower dose. As a result, we remain cautious about whether toxicity occurs in neurons at 0.5 μM. Nonetheless, even if selective cell elimination was occurring undetected, NAV 0.5 μM did not target cells in a way that is consistent with the elimination of cells expressing low LaminB1 or low nuclear HMGB1.

### Senolytic cocktail treatment with DQ does not provide evidence of selective killing of senescent cells in Aβ *in vitro* model of AD

Dasatinib acts on tyrosine kinases, including Src family kinases and Bcr-Abl^51,52^. In contrast, Quercetin interferes with anti-apoptotic pathways such as Bcl-2 and AKT survival signaling, among other targets^53,54^. The combination of Dasatinib and Quercetin in the DQ formulation was strategically developed to leverage the senolytic properties of both compounds, enabling the targeted elimination of a broader spectrum of senescent cells al lower doses^55^. DQ has been reported to ameliorate disease in a mouse model of AD-like tau pathology by selectively killing mouse senescent-like neurons whilst sparing glial cells^14^. However, probably due to the use of different transgenic mouse models, others did not report senescent neuron elimination by DQ^15^.

Experiments with Dasatinib (4 μM) and Quercetin (30 μM) followed the same methodology as those conducted for NK92 and NAV (Fig. 3a). Visual inspection did not reveal obvious pro-apoptotic effects of DQ at these doses, either in the original aCasp3 channel (Fig. 7a) or after signal enhancement in the aCasp3* channel (Sup. Fig. 7a). Similarly, MAP2 and GFAP signals remained unaffected (Sup. Fig. 7b).

**Figure 7.**
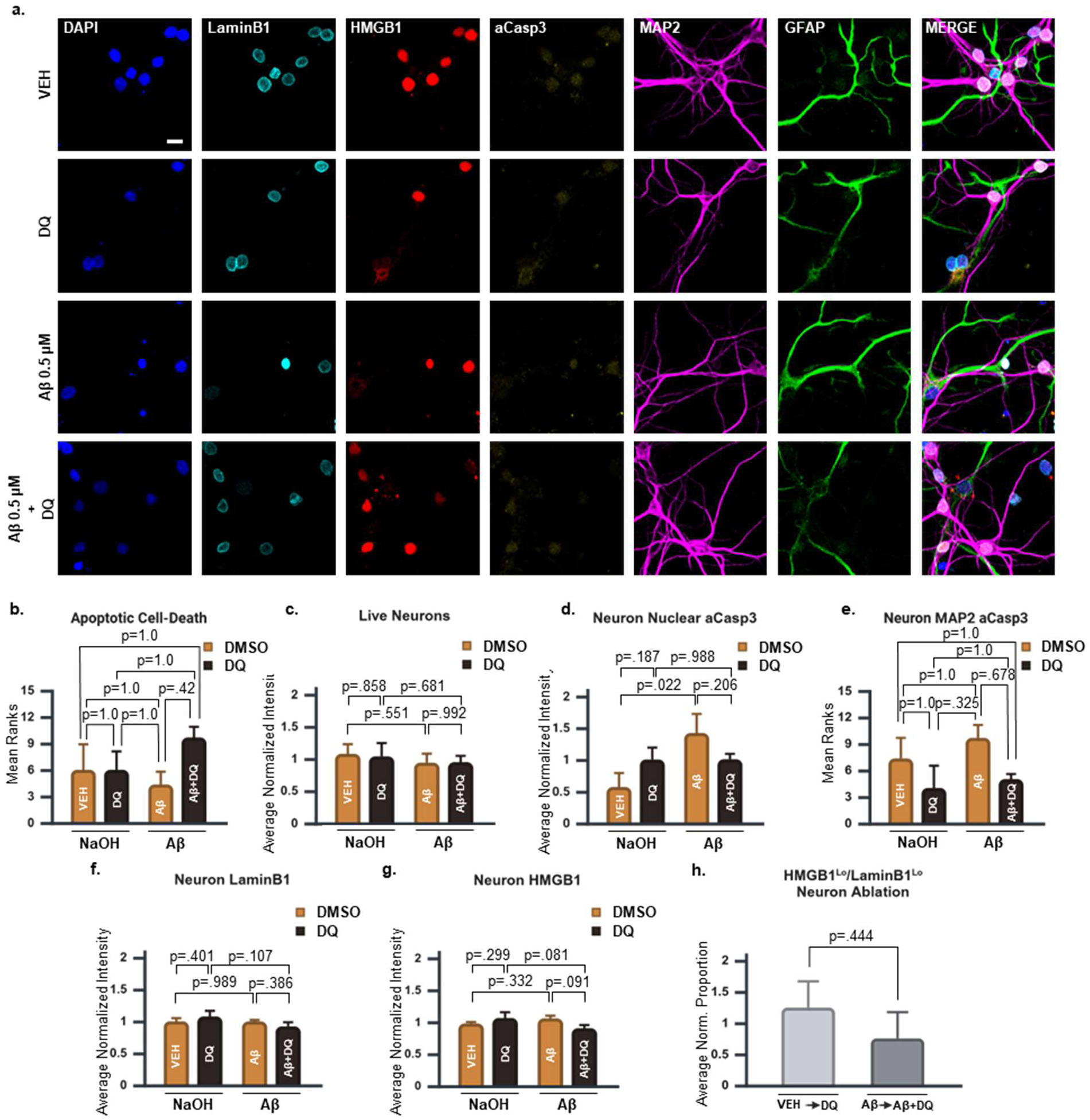
Assessment of Dasatinib 4 μM + Quercetin 30 μM senolytic cocktail on human primary neurons. (**a**) Representative linearly unmixed confocal spectral scans of all treatment conditions of DQ experiments. 40x objective with 1x zoom. Scale bar = 10 µm. (b) Mean ranks of apoptotic cell death in Aβ x DQ treatment groups, as quantified in Sup. Fig. 5e. (**c, d**) Aβ x DQ simple main effects on (c) live neuron count and (**d**) nuclear aCasp3. (**e**) Mean ranks of aCasp3 signal within MAP2 processes in Aβ x DQ treatment groups. (**f, g**) Aβ x DQ simple main effects on (**f**) LaminB1 and (**g**) nuclear HMGB1 signal. (**h**) Comparison of HMGB1^Lo^/LaminB1^Lo^ neuronal loss for VEH◊DQ and Aβ◊Aβ+DQ, as quantified in Sup. Fig 3e. (**b, e**) Significance, *p* < 0.05. (**c, d, f, g**) Significance, *p* < 0.025. (**h**) Significance, *p* < 0.0167. A total of 12313 neurons were analyzed. 3x replicates. Graphs represent mean ± s.e.m.

A non-parametric analysis revealed no statistically significant differences between treatments on apoptotic cell death (*p* = 0.319; Fig. 7b; Table 5a). There were also no statistically significant differences in the number of viable neurons analyzed across treatment conditions (Fig. 7c; Table 5b). The analysis of nuclear aCasp3 in neurons showed a trend toward significance for the interaction effect (*p* = 0.082; Table 5c), but condition-specific analysis only revealed significantly increased nuclear aCasp3 signal in Aβ-treated neurons relative to VEH that did not involve DQ (*p* = 0.022; Fig. 7d; Table 5c). Interestingly, somatodendritic aCasp3 presented a pattern more consistent with neuroprotective rather than toxic effects of DQ (Fig. 7e), but this was not ratified by a non-parametric Kruskal-Wallis H test (*p* = 0.218; Fig. 7e; Table 5d).

For analysis of cellular senescence biomarkers, LaminB1 analysis showed no significant differences that would be consistent with the elimination of specific neuron populations (Fig. 7f; Table 5e). The interaction effect between Aβ and DQ on nuclear HMGB1 did reveal a non-significant trend (*p* = 0.064; Table 5f), but subgroup analysis did not reveal significant differences or trends indicative of a biologically relevant senolytic effect (Fig. 7g). Likewise, no significant changes were observed in HMGB1^Lo^/LaminB1^Lo^ neurons when comparing the effects of DQ treatment in healthy vs. Aβ-treated neurons, providing no evidence of selective elimination (*p* = 0.444; Fig. 7h; Table 5g; Sup. Fig. 7c).

As per astrocytes, a reduction in the number of analyzed cells was associated with DQ treatment irrespective of Aβ exposure (*p* = 0.022; Fig. 8a; Table 5h), indicating non-selective loss of viability. Unexpectedly, condition-specific analysis suggested a non-significant trend toward reduced viability when comparing VEH to DQ-treated groups (*p* = 0.034), an effect not observed when comparing Aβ to Aβ+DQ groups (*p* = 0.183; Fig. 8b; Table 5h). This would reflect preferential killing of VEH-treated astrocytes, but these simple main effects should be cautiously considered given the absence of a significant interaction or trend (*p* = 0.461; Table 5h).

**Figure 8.**
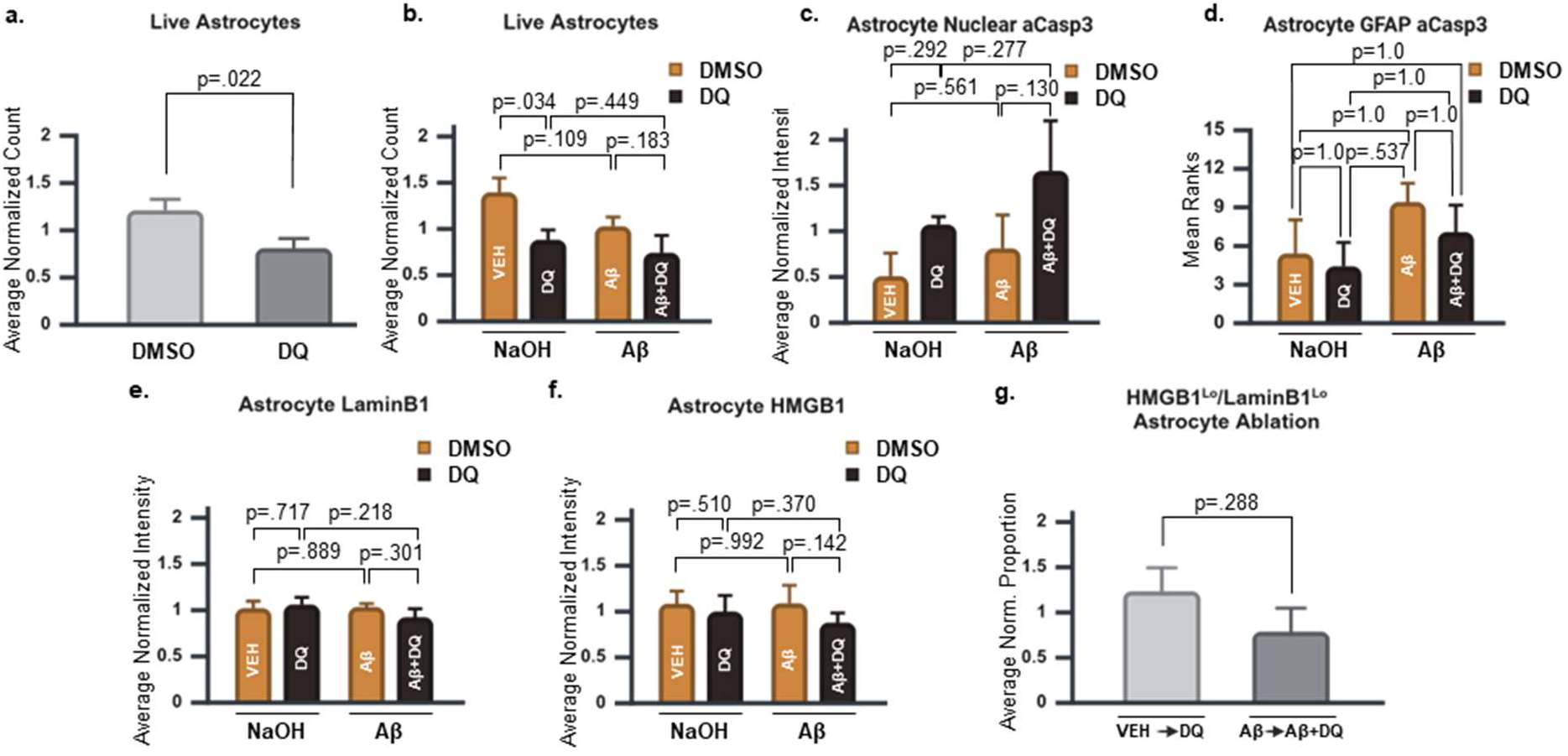
Assessment of Dasatinib 4 μM + Quercetin 30 μM senolytic cocktail on human primary astrocytes. (**a**) DQ main effects and (**b**) Aβ x DQ simple main effects on live astrocyte count. (**c**) Aβ x DQ simple main effects on nuclear aCasp3. (**d**) Mean ranks of aCasp3 signal within GFAP processes in Aβ x DQ treatment groups. (**e, f**) Aβ x DQ simple main effects on (**e**) LaminB1, and (**f**) nuclear HMGB1 signal. (**g**) Comparison of HMGB1^Lo^/LaminB1^Lo^ astrocyte loss for VEH◊DQ and Aβ◊Aβ+DQ, as quantified in Sup. Fig. 3e. (**a, d**) Significance, *p* < 0.05. (**b, c, e, f**) Significance, *p* < 0.025. (**g**) Significance, *p* < 0.0167. 3x replicates. A total of 2658 astrocytes were analyzed. Graphs represent mean ± s.e.m.

For nuclear aCasp3, a non-significant trend was observed for DQ main effects (*p* = 0.082; Table 5i). The bat chart reflected a pattern consistent with the selective elimination of Aβ-treated astrocytes, but this was not supported by significant effects in the treatment-specific analysis (Fig. 8c; Table 5i). Likewise, a non-parametric ANOVA assessing aCasp3 within GFAP-positive processes did not detect statistically significant differences (*p* = 0.347; Fig. 8d; Table 5j). The analysis of LaminB1 (Fig. 8e; Table 5k), nuclear HMGB1 (Fig. 8f; Table 5l), and HMGB1^Lo^/LaminB1^Lo^ astrocytes (Fig. 8g; Sup. Fig. 8; Table 5m) also did not show significant differences, arguing against selective killing of astrocytes by DQ.

The absence of DQ effects on neurons and astrocytes rules out senolytic activity as assessed, as well as non-selective toxicity to neurons. While our analysis suggests that DQ may induce mild, non-selective toxicity in astrocytes, statistically significant results were inconsistent across metrics, indicating that any effect—if present—is likely less pronounced than that observed with NK92 co-culture.

## Discussion

We describe a straightforward method for culturing primary human neurons and astrocytes from commercially available cryopreserved cell preparations. These cultures can be maintained well beyond 100 DIV, with GFAP-negative neurons expressing NeuN, MAP2, tau, and Ankyrin-G. By 28–43 DIV, the expression of synaptic maturation markers establishes a well-defined timeline for modeling CNS diseases with adult-onset characteristics.

### NAV, DQ, and NK92 cells: Insights into senolytic activity and selectivity

We utilized these cultures to establish an *in vitro* model of AD pathology, incorporating Aβ treatments to investigate senolytic selectivity as a proof of principle for their potential application in AD therapeutic testing.

NK92 co-culture was the only regimen in which the metrics for viability and the analysis of putative senescence markers LaminB1 and HMGB1 collectively supported the targeted elimination of senescent-like neurons and astrocytes treated with Aβ. However, elimination was partially selective at best, based on ample evidence of toxicity to neurons and astrocytes in the absence of Aβ. This aligns with previous findings on primary NK cytotoxicity on astrocytes^56^. The off-target cytotoxicity observed in this study raises concerns that CNS cells may exhibit cross-reactivity with emerging immune-based senolytic therapies designed to leverage NK signaling for senescent cell clearance^57,58^. Further studies are required to assess this potential risk.

Previous studies in mice and non-human primates have not reported NAV as neurotoxic^13,15,59^, but our results demonstrate that NAV can kill human primary neurons in a concentration-dependent manner *in vitro*. Since NAV targets the Bcl-2 family of anti-apoptotic proteins^18,48–50^, our results underscore that there may be significant risks associated with NAV and related compounds targeting this family of proteins. Future research should aim to determine whether specific targets within the Bcl-2 family can be safely leveraged to mitigate potential neurotoxicity.

In contrast, DQ did not exhibit overt neurotoxicity, even at higher concentrations commonly used *in vitro*^18^. While non-selective astrocyte toxicity was suggested by total cell counts, aCasp3-based viability metrics were inconsistent, indicating that any observed effect was likely moderate. No clear senolytic activity was detected based on LaminB1 or HMGB1, diverging from previous reports in transgenic mouse models where DQ selectively eliminated senescent-like neurons or OPCs^14,17^. However, the emergence of deleterious senescent-like neurons in previous mouse studies was attributed to tau pathology^14^, which may not be present in our cultures. Although previous studies on mouse OPCs used mouse models of Aβ pathology comparable to our in vitro experiments^17^, we neither studied nor assessed the presence of OPCs in our cultures. Consequently, our results with DQ are consistent with prior findings in mice.

While experiments with NK92 co-culture were consistent with the elimination of senescent cell populations and NAV experiments were not, neither regimen is arguably safe to target senescent cell populations that represent a small percentage of the total cell population. For example, when the senescent population represents 1% of the total population, a realistic 76% on-target and 25% off-target toxicity^18^ would theoretically result in over 3250 healthy non-senescent cells killed off-target for every 100 senescent cells killed on-target. Even if NK92 cells exhibit preferential cytotoxicity, our findings suggest their risk profile is comparable to the lack of selectivity seen with NAV at high concentrations. Given its lack of neurotoxicity, DQ is the only intervention that appears to hold promise as a truly selective senolytic. Future studies incorporating tau pathology in our model should determine whether DQ selectively eliminates senescent cells as reported in mice^14^.

## Limitations

While our findings provide valuable insights into the effects of NAV, DQ, and NK92 cells, they must be interpreted cautiously due to several limitations. We did not detect a distinct senescent-like population when comparing control and Aβ-treated cells. Aβ induces a wide range of cellular and molecular toxicities unrelated to cellular senescence^60^, which may have obscured small senescent-like populations that were revealed only under NK92 co-culture. Future studies should incorporate additional senescence markers, while acknowledging the known challenges of identifying senescent neurons^61,62^. Another limitation is that NK cells eliminate both “stressed” and senescent cells through NKG2D receptor activation^34,63^. Since Aβ is a known cellular stressor^60^, NK92 cells may have targeted stressed cells rather than or in addition to senescent cells. Additionally, NK92 cells possess a simplified receptor repertoire^64^ and our cultures lack additional immune cells required to regulate NK cytotoxicity *in vivo*. Future studies should assess NK cytotoxicity in a more physiologically relevant context. Finally, astrocyte assessments may be confounded by the use of GFAP, a marker linked to astrocyte reactivity rather than astrocyte identity^65^, emphasizing the need for additional markers to facilitate a more refined analysis.

## Conclusions

We developed an accessible method for culturing long-term primary human neurons and astrocytes using commercially available cryopreserved cell preparations. This system is straightforward to establish, requiring only basic cell culture skills, and can be readily adapted for studying a range of CNS models and therapeutic interventions. The robustness of our human *in vitro* AD model is demonstrated by its ability to differentiate the effects of NK92 cells, NAV, and DQ, aligning with their distinct mechanisms of action^18^. These findings emphasize the importance of human-specific models in refining therapeutic strategies for neurodegenerative diseases prior to their advancement to clinical applications.

## Acknowledgements

This work is supported by the Larry L. Hillblom Foundation. We thank Dr. Mark Mattson for his insights and recommendations on human primary neuron and astrocyte cultures. We thank Dr. Peter John Darlington for his insights and recommendations in NK cell co-cultures. We thank Dr. Shankar Chinta and Dr. Manish Chamoli for assistance in manuscript editing and formatting. We thank Tori Lazerson, Varun Sridhar, and Alexandra Steinberg for their work on cell cultures. We thank Allison Bischoff for insightful discussions. The figures were prepared using BioRender.

## Author Contributions

CCW, EW, SL, and CJS performed the experiments. Transductions were performed by EW and SL. Immunocytochemistry and confocal imaging were done by CCW, EW, SL, and CJS. Image processing was performed by CCW, EW, SL, and NJB. CCW performed statistical analysis. JKA and CCW supervised and guided the project. CCW wrote the manuscript, with input from all other authors.

## Materials and Methods

### Mixed human primary neuron and astrocyte cultures

To prepare Ibidi “ibiTreat” 8-well chamber slides (Ibidi, 80826) for neuronal cultures, each well was coated with 200 µL of a 1:1 solution of 1x Dulbecco’s phosphate-buffered saline (VWR, VWRL0119-0500) and Poly-D-Lysine (PDL; ThermoFisher, A3890401). The slides were placed in Petri dishes, serving as secondary containers to minimize contamination risk, and incubated overnight at 37 °C in a sterile humidified atmosphere. The following day, the wells were washed by aspirating the DPBS/PDL solution and immediately adding 300 µL of 1x DPBS to prevent drying. This washing process was repeated twice, waiting at least 2 hours between washes. The culture media was prepared before cell plating and consisted of Neurobasal Plus Medium (ThermoFisher, A3582901), 50x B-27 Plus Supplement (ThermoFisher, A3582801), 100x GlutaMAX Supplement (ThermoFisher, 35050061), and 100x Penicillin-Streptomycin solution (ThermoFisher, 15070063). Filter sterilization of the solution with a 0.22 µm syringe filter (VWR, 76479-024) prior to adding the B-27 Plus Supplement had no observable effects on cell viability. Before plating, the wells of the slides were washed with pre-warmed culture media and 240 µL of media was added to each well to accommodate approximately a 60 µL cell suspension, ensuring a final volume around 300 µL. The prepared slides were equilibrated in a humidified incubator at 37 °C with 5% CO₂ and 3-5% O₂ for at least 30 minutes before cell plating. O₂ levels at 3-5% were used to mimic physiological oxygen levels for cellular senescence experiments, but the cells may be cultured at the more conventional 20%. Vials with human fetal brain tissue between 17 and 19 gestational weeks were provided by Sciencell (Sciencell, 1520). Each cryovial (∼1 x 10⁶ cells) was briefly opened/closed to release pressure and thawed rapidly in a 37 °C water bath for approximately 2 minutes until a small ice crystal remained. The vial was then cleaned with ethanol and the cell suspension was gently tapped to dislodge cells adhering to the cap. A 50 mL centrifuge tube pre-washed with plating media was prepared to prevent cell adhesion to the tube walls. Similarly, pipette tips were primed with media before aspirating cell suspensions to minimize cell loss during handling. The cells were carefully transferred to the pre-washed tube using primed tips, with the vial cap also aspirated to recover any remaining cells. One milliliter of pre-warmed media was added dropwise to the suspension to avoid osmotic shock, followed by the gentle addition of 2 mL of media to achieve a final concentration of ∼2.5 x 10⁵ cells/mL. The cell suspension was gently pipetted to dissociate clumps and cell density confirmed using a hemocytometer. Cells were plated onto the equilibrated Ibidi slides at a density of ∼1–1.5 x 10⁴ cells/cm², resulting in a total well volume of 300 µL and placed into humidified incubator at 37 °C with 5% CO₂ and 3-5% O₂. Completely replacing the media after plating to remove residual cryopreservants significantly reduced cell viability. Omitting the media replacement step and leaving the cells completely undisturbed for 48 hours post-plating substantially improved culture health. Subsequent media changes were performed every 2 days whenever feasible, replacing only one-third of the media with fresh culture media lacking Penicillin-Streptomycin. Retaining sufficient conditioned media during these changes was essential to support the cells. Adjustments to the media volume were made to account for evaporation, ensuring the total well volume remained consistent with the initial plating volume. At the specified plating density, antimitotics were not required as astrocyte growth did not adversely affect neuronal viability or the overall health of the cultures. While some slight variations in cell density and volume can occur, both the overall cell-to-volume and cell-to-area ratios described herein were found to be important for the long-term viability of the cultures.

### NK92 culture and co-culture

NK92 cells (ATCC, CRL-2407) were cultured following the vendor’s instructions. Complete growth media was prepared using alpha MEM media (Gibco, 12561072) supplemented with 12.5% non-heat-inactivated fetal bovine serum (VWR, 45000-73), 12.5% heat-inactivated horse serum (ThermoFisher, 26050088), 0.02 mM folic acid (Sigma, F8758-5G), 0.2 mM myo-Inositol (Sigma, I7508-50G), 0.1 mM beta-mercaptoethanol (ThermoFisher, 21985023), 100 U/mL Penicillin-Streptomycin solution (ThermoFisher, 15070063), and 100 U/mL human recombinant IL-2 (R&D Systems, 202-IL). The prepared media was equilibrated in T25 flasks (Corning) in humidified incubator at 37 °C with 5% CO₂ and 20% O₂. Thawed NK92 cells were centrifuged at 800 x g for 5 minutes, resuspended in pre-warmed complete growth media, and transferred to pre-equilibrated T25 flasks at an initial density of 2–4 x 10⁵ cells/mL. Cultures were maintained in a humidified incubator and split every 2–3 days, with the culture medium refreshed to sustain optimal cell density and viability. For co-culture experiments, NK92 cells were harvested, pelleted at 340 x g for 7 minutes, and resuspended in neurobasal medium. The cells were subsequently added to 42 DIV mixed primary human neuron and astrocyte cultures at an effector-to-target (E:T) ratio of 10:1 and co-cultured for 24 hours (Figures 3 & 4) or 20 hours (Videos 10-13).

### Live Imaging

Cells were cultured in Ibidi 8-well chamber slides with polymer coverslip bottoms coated with PDL. Live-cell imaging was conducted using a Zeiss LSM 780 confocal microscope (Carl Zeiss Microscopy GmbH, Jena, Germany) with 20x, 40x, or 63x objectives. Imaging was performed in a sealed chamber maintained at 37°C and 5% CO₂. Images and videos were captured using ZEISS ZEN Microscopy Software (RRID:SCR_013672). Details regarding objectives, time intervals between frames, and experiment durations are provided in the video legends.

### Cytoskeleton visualization

The cytoskeleton was visualized by transducing Tubulin tagged with red fluorescent protein (Tubulin-RFP) using the CellLight Tubulin-RFP, BacMam 2.0 reagent (ThermoFisher, C10503), following the manufacturer’s instructions. Briefly, the CellLight reagent was added to wells containing 15,000 cells at a multiplicity of infection of 70 particles per cell and incubated for 16 hours. Subsequently, cells were observed under a 40x objective on the Zeiss LSM 780 confocal microscope.

### Aβ & Senolytic Treatments

Aβ peptide preparations enriched in oligomers were generated using a Beta Amyloid (1-42) aggregation kit (rPeptide, A-1170-2). Lyophilized Aβ peptides were reconstituted in either 5 mM Tris or 10 mM NaOH at a concentration of 1 mg/mL, then diluted with HPLC-grade water and Tris-buffered saline (TBS) to create a 100 µM stock solution. To enrich for oligomers, the stock solution was incubated at 37°C for 3 hours, aliquoted, and stored at −20°C until use. Primary human neuronal-astrocytic cultures were treated with Aβ preparations or their vehicle at 28 DIV at a final concentration of 0.5 µM. The senolytics Navitoclax (Selleckchem, S1001), Dasatinib (Selleckchem, S1021), and Quercetin (Selleckchem, S2391) were prepared according to the vendor’s instructions. NAV (0.5, 4, or 8 µM), Dasatinib (4 µM) plus Quercetin (30 µM), or a DMSO vehicle solution (final DMSO concentration < 0.1%), were added to the cultures at 42 DIV. After 24 hours of treatment, the cultures were collected for ICC.

### Antibodies

Primary antibodies were used at the following dilutions: The Lamin B1 rabbit polyclonal antibody (Abcam, ab16048), 1:1000; HMGB1 (1F3) IgG2a mouse monoclonal antibody (Abcam, ab190377), 1:1000; cleaved (active) Caspase 3 (Asp175) (5A1E) rabbit monoclonal antibody (Cell Signaling, 9664), 1:250; GFAP (2.2B10) rat IgG2a monoclonal antibody (Invitrogen, 13-0300), 1:1500; GFAP rabbit polyclonal antibody (Abcam, ab7620), 1:1000; MAP2 chicken polyclonal antibody (Abcam, ab5392), 1:2500; NeuN (1B7) IgG2b mouse monoclonal antibody (Abcam, ab104224), 1:500-1:1000; tau (Tau-5) IgG1 mouse monoclonal antibody (Abcam, ab80579), 1:200; 3R tau (8E6/C11) mouse monoclonal antibody (Sigma, 05-803), 1:100; tau (MN423) chimeric rabbit antibody (Absolute Antibody, Ab02389-23.0), 1:100; tau (MC1) IgG1 mouse monoclonal antibody (a kind gift from Dr. P. Davies), 1:50; ankyrin-G (N106/36) IgG2a mouse monoclonal antibody (Sigma, MABN466), 1:50; PSD-95 (6G6-1C9) IgG2a mouse monoclonal antibody (ThermoFisher, MA1-045), 1:500; Homer-1 IgG1 mouse monoclonal antibody (SYSY, 160 011), 1:200; Synapsin-1 (A6442) rabbit polyclonal antibody (ThermoFisher, A6442), 1:250. Secondary antibodies were used at the following dilutions: Goat anti-Rabbit IgG Alexa Fluor 488 (Invitrogen, A-11008), goat anti-Rabbit IgG Alexa Fluor 488 (Invitrogen, A-11035), goat anti-Rabbit IgG Alexa Fluor 647 (Invitrogen, A-32733), goat anti-Mouse IgG Alexa Fluor 546 (Invitrogen, A-11030), and goat anti-Mouse IgG1 Alexa Fluor 594 (Invitrogen, A-21125), 1:500; goat anti-Mouse IgG2b Alexa Fluor 546 (Invitrogen, A-21143), goat anti-Rabbit IgG Alexa Fluor 594 (Invitrogen, A-32740), goat anti-Rat IgG Alexa Fluor 633 (Invitrogen, A-21094), and goat anti-Mouse IgG1 Alexa Fluor 488 (Invitrogen, A-21121), 1:1000; goat anti-Chicken IgY Alexa Fluor 647 (Invitrogen, A-21449) and goat anti-Mouse IgG Alexa Fluor 488 (Invitrogen, A-11001), 1:1000-1:2000.

### Immunocytochemistry (ICC)

ICC and confocal imaging were performed on the microscopy-compatible Ibidi slides to ensure minimal disruption to cellular morphology. Cell cultures were fixed by progressively replacing the culture media with 4% paraformaldehyde (PFA; Santa Cruz, 3052589-4) and incubated with PFA at room temperature (RT) for 15 minutes, followed by three washes with ice-cold PBS for 10 minutes each. Permeabilization was performed using PBS containing 0.05% Triton x-100 (Northwest Scientific, JTB/x198-07) (PBTx) for 30 minutes. After permeabilization, cultures were incubated for 30 minutes at RT in PBTx supplemented with 10% normal goat serum (NGS; Jackson ImmunoResearch, 005-000-121) to block nonspecific antibody binding. Primary antibodies were diluted in PBS containing 1% NGS and applied to the cultures for 1 hour at RT. This was followed by four 10-minute washes with PBS to remove unbound primary antibodies. Secondary antibodies, diluted in PBS with 1% NGS, were applied for 1 hour at RT, followed by four additional 10-minute washes with PBS. For ICC involving repeated use of isotype-specific antibodies, a sequential staining protocol was employed. The first round of primary and secondary antibody incubations was performed on the first day. Following an overnight wash to remove residual antibodies, a second round of primary and secondary antibody incubations was carried out on the next day. DNA labeling was performed by incubating the cultures in PBS containing 1 μg/mL 4′,6-diamidino-2-phenylindole (DAPI; Invitrogen, D1306) for 3 minutes. After three final washes with PBS, the PBS was entirely replaced with pure glycerol (Invitrogen, 15514-011) to preserve the stained cells and minimize refractive index changes during imaging. Confocal imaging was performed using a Zeiss LSM 980 for cell culture characterization and Zeiss LSM 780 confocal microscopes (Carl Zeiss Microscopy GmbH, Jena, Germany) for cell culture characterization and senolytic regimen studies.

### Image Generation and Processing

Maximum intensity projection (MIP) images and time-lapse videos were created using ZEISS ZEN Microscopy Software (RRID:SCR_013672). IMARIS software (RRID:SCR_007370) was used for 3D visualization, video generation, and processing of 3D confocal images. The algorithms used to generate the ROIs were optimized for each replicate and standardized across treatments within the same replicate to eliminate user bias. **Synapse Characterization.** ROI for Synapsin, Homer-1, and PSD95 were defined using the spot function in IMARIS. The sum voxel intensity within all ROIs for each DIV and replicate in each channel was calculated and used for statistical analysis of total protein levels. To quantify colocalization between Synapsin-Homer-1 and Synapsin-PSD95, spots generated for the above-described analysis were used to generate new channels with the synaptic markers. The IMARIS colocalization function was applied to generate a new channel containing colocalized voxels of the masked channels. The spot function was then used on the channel to generate ROI on colocalized puncta. The total number of ROIs for colocalized puncta was calculated for each DIV and replicate and used for statistical analysis.

### Viability, LaminB1, and HMGB1 Analysis for NK92, NAV 0.5 µM, and DQ Experiments

Nuclear ROIs were defined using the surface function applied to DAPI-stained nuclei. User-defined algorithms were utilized to classify cells as live or dead based on nuclear characteristics. Additional user-defined and machine learning algorithms were employed to further classify live cell nuclei ROI as belonging to neurons or astrocytes, with manual inspection performed to validate the classifications. The processed nuclear ROI data were exported as CSV files and imported into FlowJo (RRID:SCR_008520) for further refinement. Within FlowJo, neurons were identified as MAP2^hi^/GFAP^lo^ and astrocytes as MAP2^lo^/GFAP^hi^ by plotting MAP2 against GFAP channels. These populations were then exported into new CSV files for subsequent statistical analyses. The total number of ROIs classified as neurons or astrocytes was used to analyze the viability of live cells. Mean HMGB1 and aCasp3 intensity values within the ROIs were used for statistical analyses of nuclear HMGB1 and aCasp3 levels. Additional steps were required to accurately assess LaminB1. Nuclear-based ROIs inherently overcapture the LaminB1 signal, a nuclear lamina protein, and average its signal with non-nuclear lamina regions within the ROI. This results in an underestimation of LaminB1 signal intensity in large nuclei relative to smaller nuclei. To address this issue, a new LaminB1 channel was created by setting LaminB1-positive voxels in the original channel to the maximum intensity (255 in an 8-bit image). Within each nuclear ROI, the sum intensity of this newly generated channel was divided by 255 to calculate the number of LaminB1-positive voxels, effectively excluding contributions from non-nuclear lamina regions. The sum signal intensity of the original LaminB1 channel was then divided by the number of LaminB1-positive voxels to obtain the true LaminB1 mean intensity for each ROI. This corrected LaminB1 mean was subsequently used for statistical analysis.

### Apoptotic Cell Death Analysis for NAV and DQ Experiments

Pyknotic nuclei were identified in the DAPI channel using user-defined algorithms, generating corresponding ROIs. An enhanced aCasp3 channel (aCasp3*) was created by increasing the relative intensity of aCasp3-positive voxels inversely proportional to signals in other channels (as shown in Sup. Fig. 5b). Noise in the aCasp3* channel was minimized using Gaussian, median, and threshold cutoff filters in IMARIS where necessary, standardizing the steps for all treatments within each replicate. This refined aCasp3* channel was used to define ROIs representing apoptotic cell remnants. These ROIs were size limited to fragment larger apoptotic remnants into smaller, discrete ROIs overlapping with pyknotic nuclei ROIs. The pyknotic and aCasp3* ROIs corresponding to the same apoptotic cell remnant were identified based on spatial proximity and visually inspected to ensure accuracy. Subsequently, these ROIs were used to mask a new channel from which final ROIs representing apoptotic cell bodies were generated (Sup. Fig. 5e). Additional ROIs were created to identify live cells, irrespective of cell type, to determine the total number of live cells in the cultures. The proportion of apoptotic cell ROIs to live cell ROIs was calculated and used for statistical analysis, providing a measure of apoptotic cell death relative to overall cell viability within the cultures.

### aCasp3 Analysis in Neuronal Processes for NK92, NAV 0.5 µM, and DQ Experiments

ROI for neuronal and astrocytic processes were defined using the surface function of IMARIS on MAP2 and GFAP channels, respectively. The sum pixel intensity of aCasp3 within each ROI was calculated and divided by the ROI volume to obtain a volume-normalized aCasp3 signal. This normalized signal was used for statistical analyses to assess cytoplasmic aCasp3.

### Volume Analysis of Neuronal and Astrocytic Processes for NAV 4 and 8 µM Experiments

To analyze neuronal and astrocytic processes, ROIs were created based on MAP2 and GFAP channels. The total volume of MAP2-positive or GFAP-positive ROIs was calculated and divided by the confocal image area to determine the volume of neuronal or astrocytic processes normalized to the image area, which was subsequently used for statistical analyses.

### Statistics

Statistical analyses were conducted using IBM SPSS Statistics (RRID:SCR_002865). Both significant results and non-significant trends are reported in the main text, while non-significant findings without trends are omitted from the text for clarity. Instead, details of all statistical tests, including assumption tests, are presented in Tables 1–5. Assumptions tests were carried out to detect extreme outliers (box plots), normality (Shapiro-Wilk test, *p* > 0.05, and Q-Q plots), and homogeneity of variances (Levene’s test, *p* > 0.05). Transformations appropriate to the type of assumption violation were attempted and assumption tests re-run. When transformed data did not meet the assumptions, non-parametric tests were run. **Synapse maturation:** One-way ANOVA or Welch’s F omnibus tests and Tukey’s or Games-Howell multiple comparisons were performed as determined by Levene’s test for homogeneity of variances. Statistical significance was set at *p* < 0.05 and trends at p < 0.1 (two-tailed). The sum voxel signal intensity within the ROIs for Synapsin, Homer-1, and PSD95 was calculated replicate by replicate for each DIV to quantify total protein levels. Colocalization between Synapsin and Homer-1, as well as Synapsin and PSD95, was assessed by calculating the total number of ROIs capturing colocalized puncta, as defined in the image processing protocol. The analysis was performed on the log10 transform on the sum signal intensity or sum number of colocalized puncta for each DIV replicate by replicate. **Senolytic regimens:** For the analysis of the effects of senolytic regimens on viable neurons, LaminB1, and nuclear HMGB1 measured by ROI based on nuclei, Mahalanobis distances were used to detect and discard multivariate outliers from the nuclei ROI data, with statistical significance assessed using a Chi-squared test (*p* < 0.01). The nuclear ROI predictive variables included voxel number, the length across the vertical axis, LaminB1 mean, HMGB1 mean, the shortest, mid, and longest distances within the ROI bounding box. After removing outliers, treatment means for cell count (Live neurons/astrocytes), nuclear aCasp3, LaminB1, and HMGB1 were calculated replicate by replicate and normalized to the average of each replicate. HMGB1^Lo^/LaminB1^Lo^ were calculated as described in Sup. Fig. 3e and normalized to the average of each replicate. For the analysis of apoptotic cell death (Sup. Fig. 5e; see image processing), MAP2/GFAP volume per area (see image processing), and aCasp3 signal within MAP2/GFAP (see image processing), treatment means were calculated replicate by replicate and normalized to the average of each replicate. For the resulting variables, assumptions tests and transformations were carried out as described above for synapse maturation with the addition of scatter plots of predicted and residual values to complement Levene’s test. A two-way ANOVA was used to assess the interaction and the main effects. For 2×2 ANOVA used in NK92, NAV 0.5 μM, and DQ, the interaction assesses whether the difference between the double vehicle and the senolytic regiment is significantly different from the difference between Aβ treatments and Aβ plus the senolytic regimen. Regardless of the presence of an interaction, hypothesis-driven simple main effects for Aβ and for the senolytic regimen were run to identify differences relevant to the study while omitting irrelevant comparisons. The simple main effects for Aβ compare the effects of adding Aβ in the absence of senolytics and, separately, in the presence of senolytics. The simple main effects for the senolytic regimen compare the effects of adding senolytic regimen in the absence of Aβ and, separately, in the presence of Aβ. Bar graphs with simple main effect *p* values are always plotted. Main effects are only plotted when they reach statistical significance in the absence of a significant interaction. Bonferroni correction was applied to correct for two simple main effects, declaring significance at *p* < 0.025 and a non-significant trend at *p* < 0.05. When two-way ANOVA assumptions were not met, a non-parametric one-way ANOVA on ranks (Kruskal-Wallis) was run followed by Bonferroni all-pairwise comparisons (*p* < 0.05). The assessment of HMGB1^Lo^/LaminB1^Lo^ populations was performed using two-tailed independent t-tests. Because the HMGB1 and LaminB1 datasets were previously analyzed in two-way ANOVAs, their reuse introduced the potential for inflated Type I error. To mitigate this, a Bonferroni correction was applied for three tests, adjusting the significance threshold to p < 0.0167. This stricter threshold accounted for multiple analyses involving the same data to control for the likelihood of false-positive results.

