## Supplementary Materials for "A Human Neuron Alzheimer’s Disease Model Reveals Barriers to Senolytic Translatability"

**Video Legends.**

**Video 1.** Transmitted light time-lapse video of cells 2 hours after plating. Note lamellipodium actively probing the environment. 63x objective with 1.5x zoom.

**Video 2.** Confocal time-lapse video of cells expressing CellLight™ Tubulin-RFP (BacMam 2.0) for visualization of the cytoskeleton. 24 hours after plating. Note RFP-positive cells actively probing the environment. 40x objective with 0.6x zoom.

**Video 3.** Confocal time-lapse video of cells expressing CellLight™ Tubulin-RFP (BacMam 2.0) for visualization of the cytoskeleton. 24 hours after plating. Note a cell migrating from the cluster of cells (top left), clearly visible at time 06:26:19.331. Note cells undergoing cytokinesis (bottom right), starting at time 02:56:57.874. 40x objective with 0.6x zoom.

**Video 4.** Confocal plus transmitted light time-lapse video of cells expressing CellLight™ Tubulin-RFP (BacMam 2.0) for visualization of the cytoskeleton. 24 hours after plating. Note cytokinesis onset of central RFP-positive cell first detected at time 04:02:22.261. 40x objective with 0.6x zoom.

**Video 5.** Confocal plus transmitted light time-lapse video of cells expressing CellLight™ Tubulin-RFP (BacMam 2.0) for visualization of the cytoskeleton. 24 hours after plating. Note the astrocyte-like morphology of the RFP-positive cell (bottom, left corner) compared to the long axon-like projection of a potential developing neuron (top, right corner). 40x objective with 0.6x zoom.

**Video 6.** Transmitted light time-lapse video of control primary neuron and astrocyte cultures from 42 to 43 DIV. 40x objective with 1x zoom.

**Video 7.** Transmitted light time-lapse video of primary neuron and astrocyte cultures co-cultured with NK92 cells from 42 to 43 DIV. 40x objective with 1x zoom.

**Video 8.** Transmitted light time-lapse video of Aβ-treated primary neuron and astrocyte cultures co-cultured with NK92 cells from 42 to 43 DIV. Aβ treatments (0.5 μM) were carried out at 28 DIV. 40x objective with 1x zoom.

**Video 9.** Video of 3D reconstructed confocal spectral scan with linear unmixing images of 43 DIV mixed human primary neuron and astrocytes co-cultured with NK92 cells stained for MAP2 (violet) to identify neurons, GFAP (green) to identify astrocytes, LaminB1 (light green) and HMGB1 (not shown) as proxies for cellular senescence, DAPI (blue) for nuclei, and aCasp3 (yellow) to identify apoptotic cells. Close up of a double GFAP and aCasp3 positive astrocyte is shown at second 48.

**Video 10.** Video of 3D reconstructed confocal spectral scan with linear unmixing images of VEH-condition 43 DIV mixed human primary neuron and astrocyte cultures stained for MAP2 (violet) to identify neurons, GFAP (green) to identify astrocytes, DAPI (blue) for nuclei, and aCasp3 (yellow) to identify apoptotic cells. Neuronal (MAP2, violet) and astrocyte (GFAP, green) processes appear normal with few aCasp3 positive apoptotic cell remnants.

**Video 11.** Video of 3D reconstructed confocal spectral scan with linear unmixing images of 43 DIV mixed human primary neuron and astrocyte cultures treated with Aβ and stained for MAP2 (violet) to identify neurons, GFAP (green) to identify astrocytes, DAPI (blue) for nuclei, and aCasp3 (yellow) to identify apoptotic cells. Neuronal (MAP2, violet) and astrocyte (GFAP, green) processes appear normal with few aCasp3 positive apoptotic cell remnants.

**Video 12.** Video of 3D reconstructed confocal spectral scan with linear unmixing images of 43 DIV mixed human primary neuron and astrocyte co-culture with NK92 cells stained for MAP2 (violet) to identify neurons, GFAP (green) to identify astrocytes, DAPI (blue) for nuclei, and aCasp3 (yellow) to identify apoptotic cells. Neuronal (MAP2, violet) and astrocyte (GFAP, green) processes strongly deteriorated with abundant aCasp3 positive apoptotic cell remnants and astrocytes, showcasing extensive off-target cytotoxicity to vehicle control cells.

**Video 13.** Video of 3D reconstructed confocal spectral scan with linear unmixing images of 43 DIV mixed human primary neuron and astrocyte treated with Aβ and co-cultured with NK92 cells stained for MAP2 (violet) to identify neurons, GFAP (green) to identify astrocytes, DAPI (blue) for nuclei, and aCasp3 (yellow) to identify apoptotic cells. Neuronal (MAP2, violet) and astrocyte (GFAP, green) processes strongly deteriorated with abundant aCasp3 positive apoptotic cell remnants and astrocytes, showcasing extensive cytotoxicity.

a.

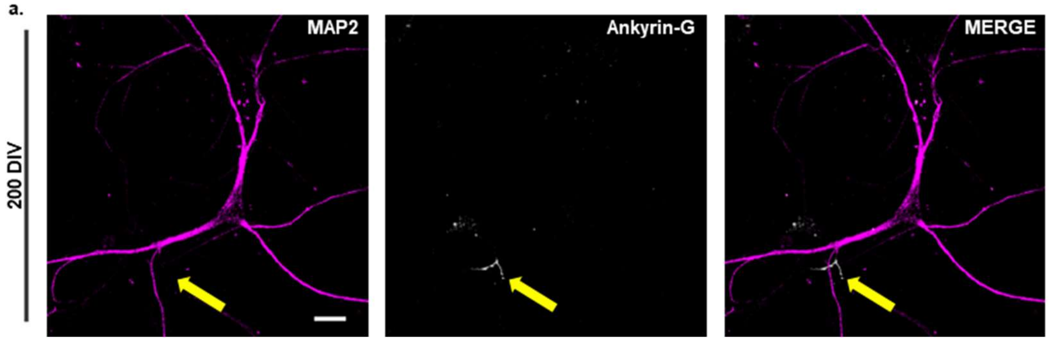


**Sup. Fig. 1a.** Representative linearly unmixed confocal spectral scans of MAP2-positive neuron at 200 DIV of culture. MAP2 labels somatodendritic neuronal processes and Ankyrin-G (yellow arrow) the axon initial segment. 40x objective with 1x zoom. Scale bar = 20 μm.

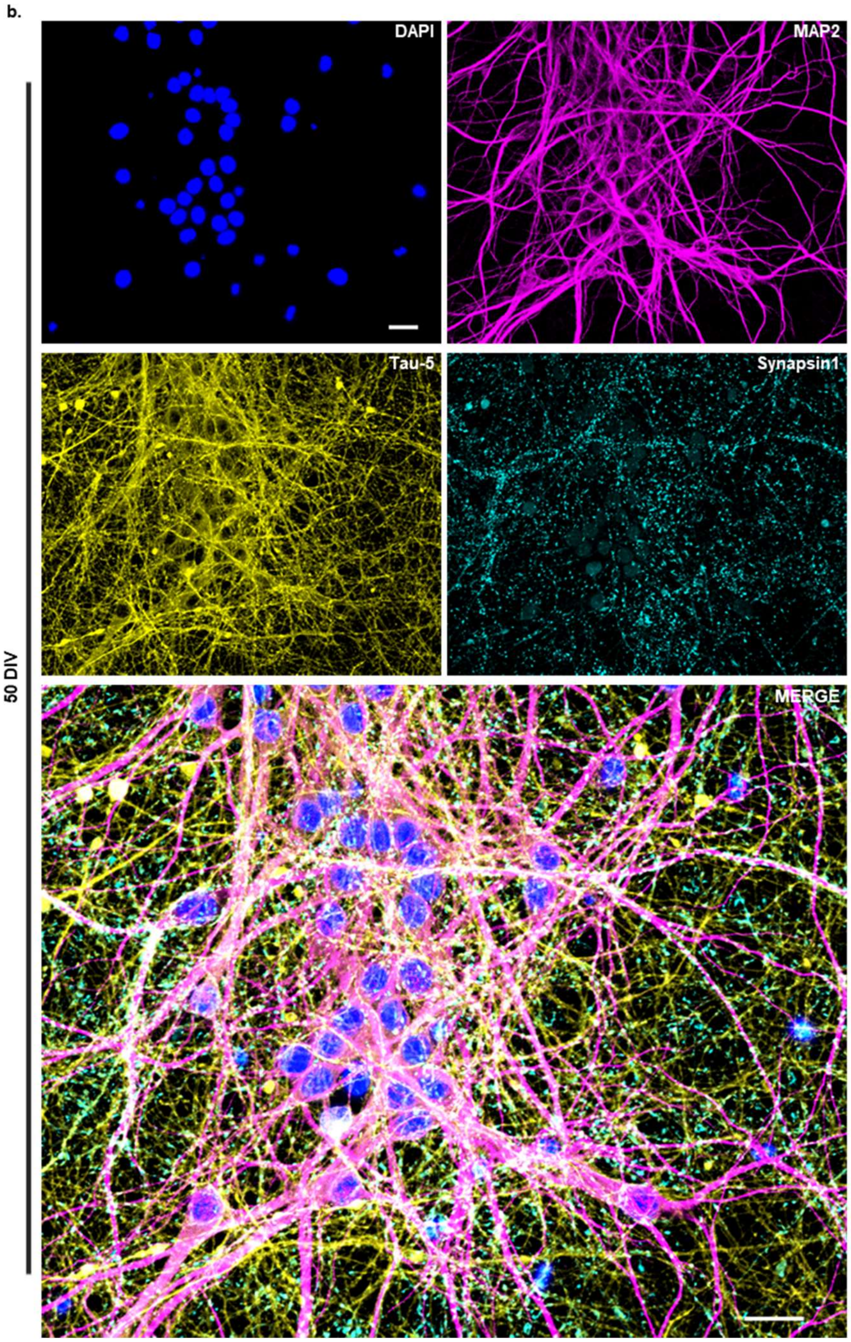


**Sup. Fig. 1b.** Confocal microscopy tile scan image of mixed primary human neuron cultures at 50 DIV. DAPI staining identifies nuclei, MAP2 labels somatodendritic neuronal processes, Tau-5 is a pan-tau axonal marker, and Synapsin1 identifies axon terminals. Merged image is enlarged. 63x objective with 0.6x zoom. Scale bar = 20 µm.

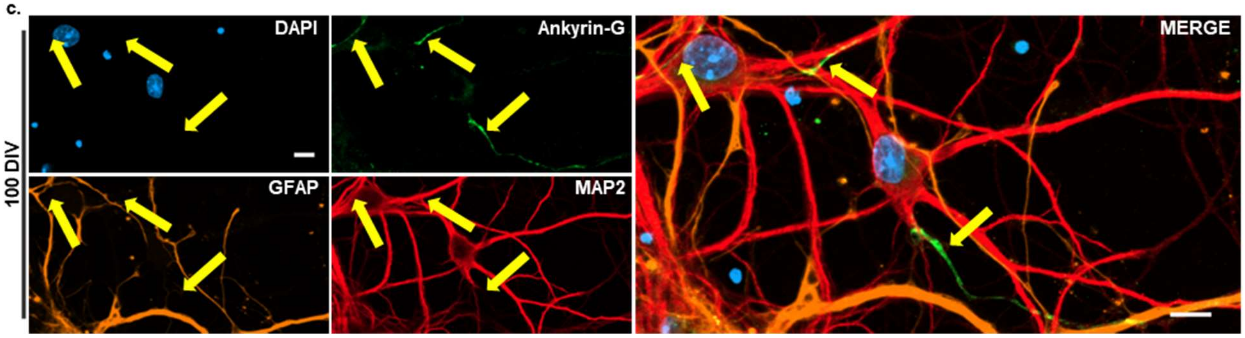


**Sup. Fig. 1c.** Super-resolution microscopy tile scan image of MAP2-positive neurons from 100 DIV cultures, showing Ankyrin-G localization at the axon initial segment (yellow arrows). DAPI staining highlights nuclei, and GFAP identifies astrocytes. 63x objective with 1.7x zoom. Scale bars = 10 µm.

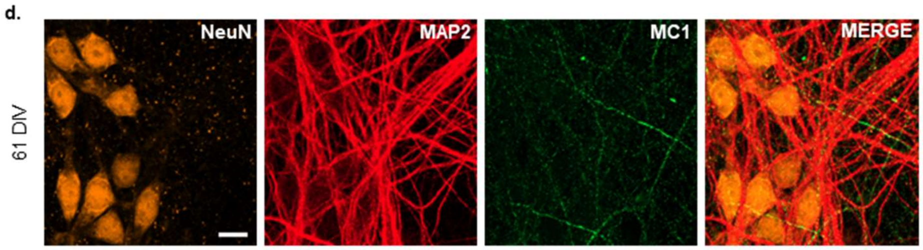


**Sup. Fig. 1d.** Confocal microscopy image of NeuN and MAP2-positive neurons from 61 DIV cultures, showing MC1 tau antibody positive immunostaining of axons. 40x objective with 2x zoom. Scale bar = 10 μm.

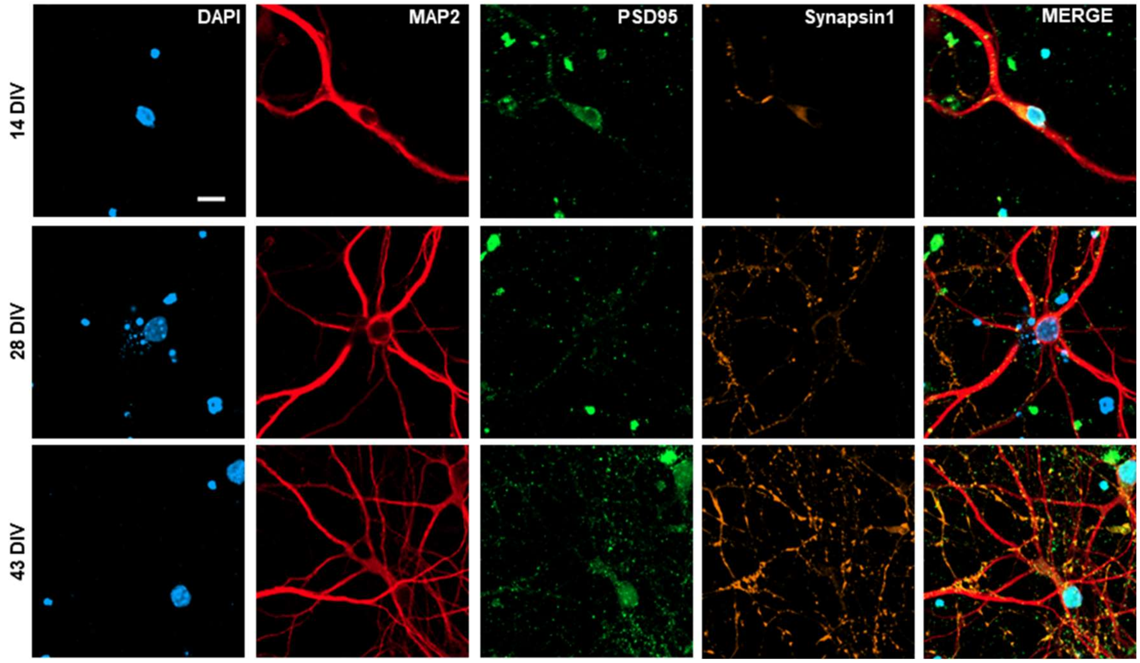


**Sup. Fig. 2.** Representative super-resolution confocal microscopy images from cultures at 14, 28, and 43 DIV. DAPI was used to identify nuclei, MAP2 to label somatodendritic neuronal processes, PSD95 as a postsynaptic marker, and synapsin as a presynaptic marker. 63x objective with 2x zoom. Scale bar = 10 µm.

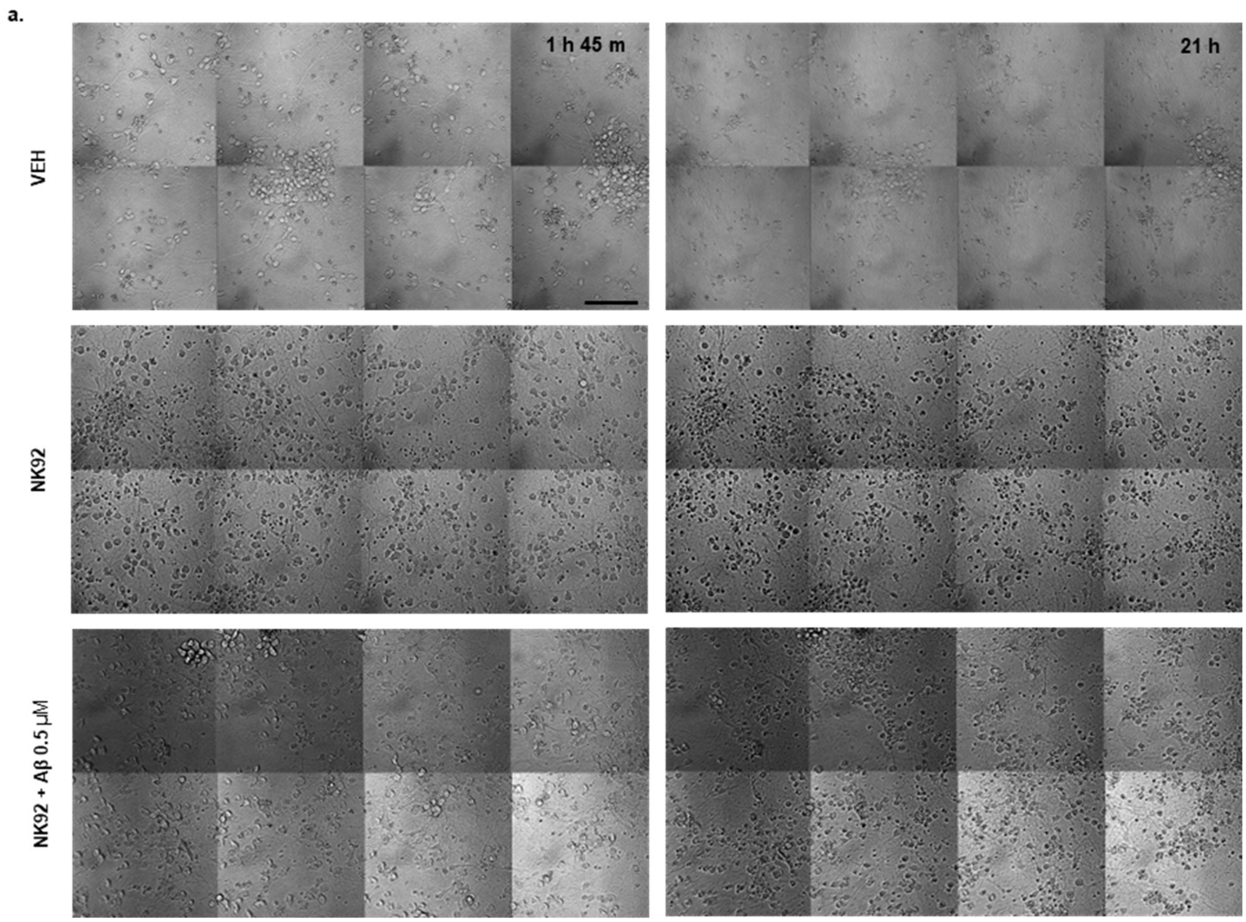


**Sup. Fig. 3a.** Stills from transmitted light tile scan images of time-lapse experiments in videos 6 (top), 7 (middle), and 8 (bottom) 1 hour and 45 minutes (left) and 21 hours (right) after adding NK92 cells to mixed human primary neuron cultures at 42 DIV. 20x objective with 1x zoom. Scale bar = 100 μm.

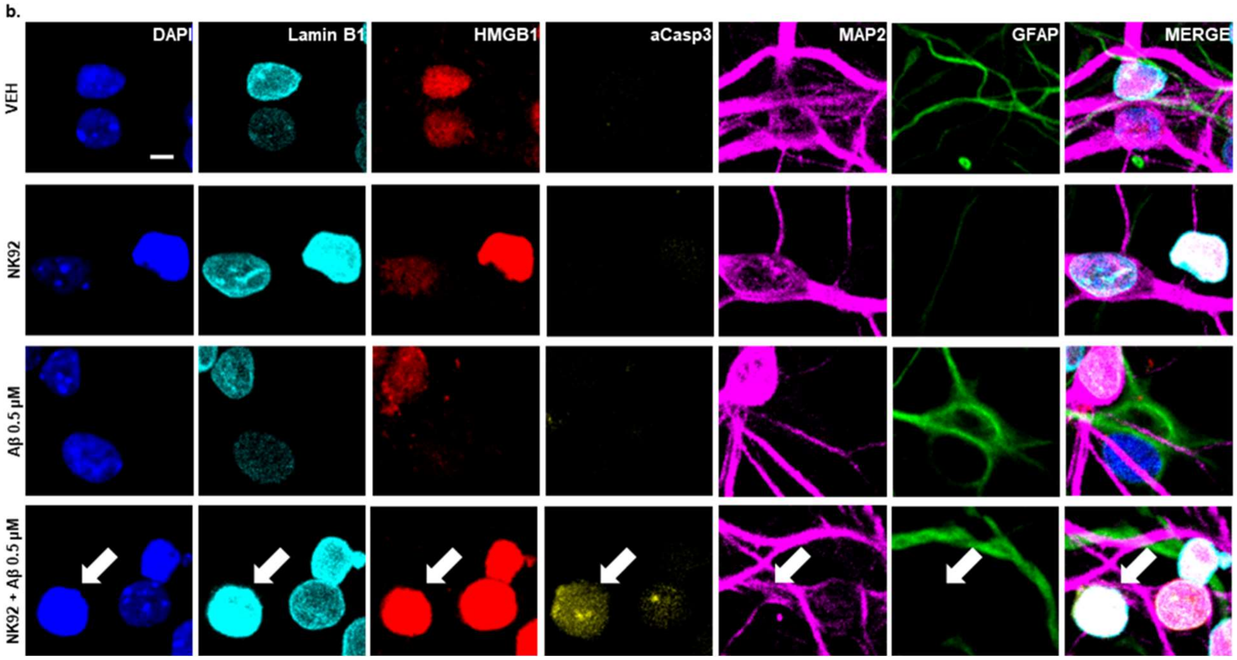


**Sup. Fig. 3b.** Representative linearly unmixed confocal spectral scans of VEH, NK92, Aβ, and Aβ + NK92 groups with ICC for DAPI, LaminB1, HMGB1, aCasp3, GFAP, MAP2 and merged images. Aβ and Aβ + NK92 images are also presented in Fig. 3b. The white arrow identifies a MAP2 and GFAP negative cell with relatively strong nuclear aCasp3 that is likely an NK92 cells. 40x objective with 1x zoom. Scale bar = 5 μm.

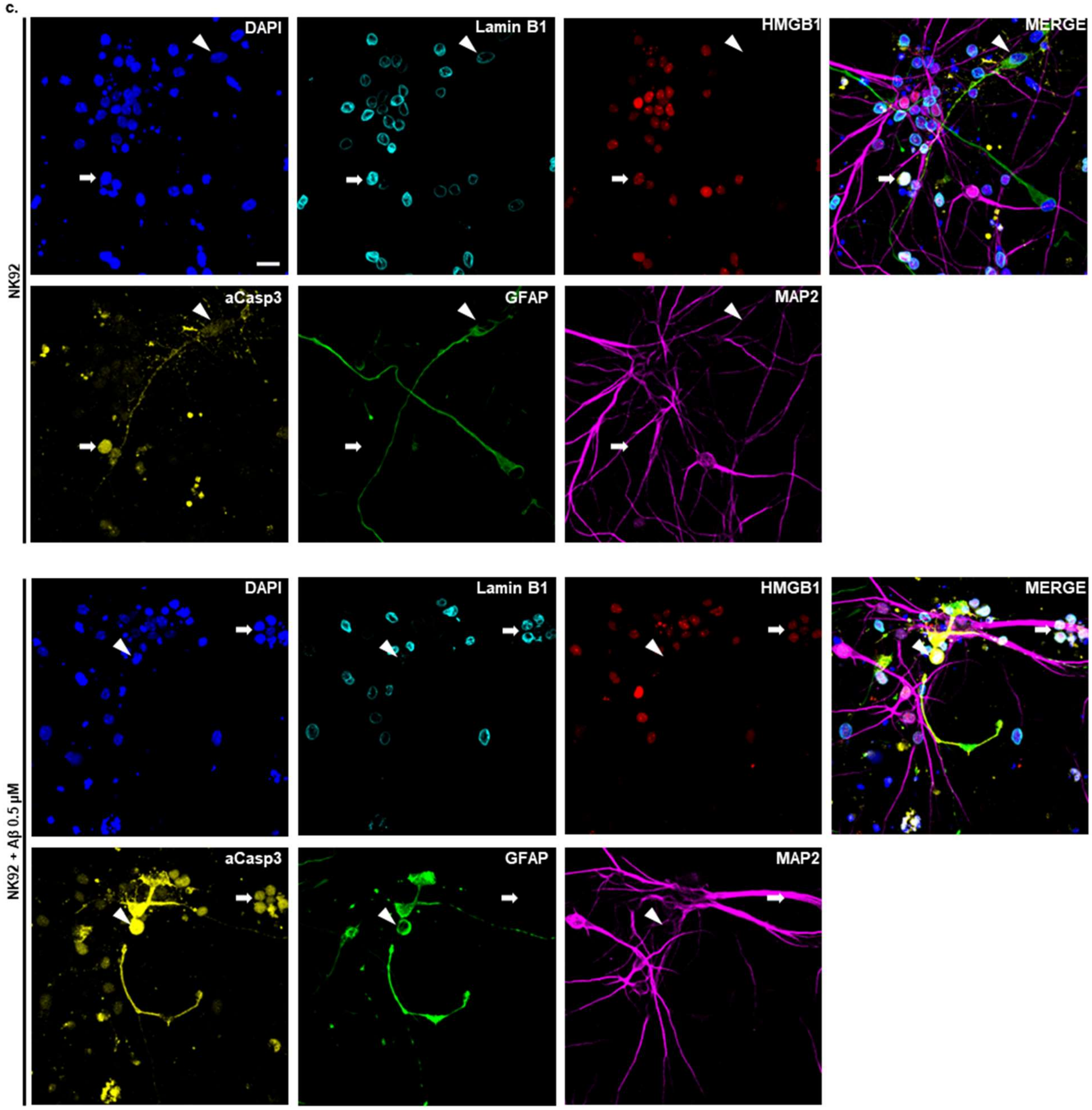


**Sup. Fig. 3c.** Representative linearly unmixed confocal spectral scans of NK92 and NK92 + Aβ 0.5 μM groups with ICC for DAPI, LaminB1, HMGB1, aCasp3, GFAP, MAP2 and merged images. For NK92 images, the white arrow identifies a MAP2 and GFAP negative cell with relatively strong nuclear aCasp3 that is likely an NK92 cell. The white arrowhead identifies a GFAP-positive live astrocyte actively undergoing apoptosis, as evidenced by positive LaminB1 signal despite the presence of aCasp3 staining. For NK92 + Aβ 0.5 μM images, the white arrow identifies a group of putative NK92 cells atop MAP2-positive processes. The white arrowhead identifies the nucleus of a GFAP-positive astrocyte undergoing apoptosis that is negative for LaminB1 and HMGB1. 40x objective with 1x zoom. Scale bar = 20 μm.

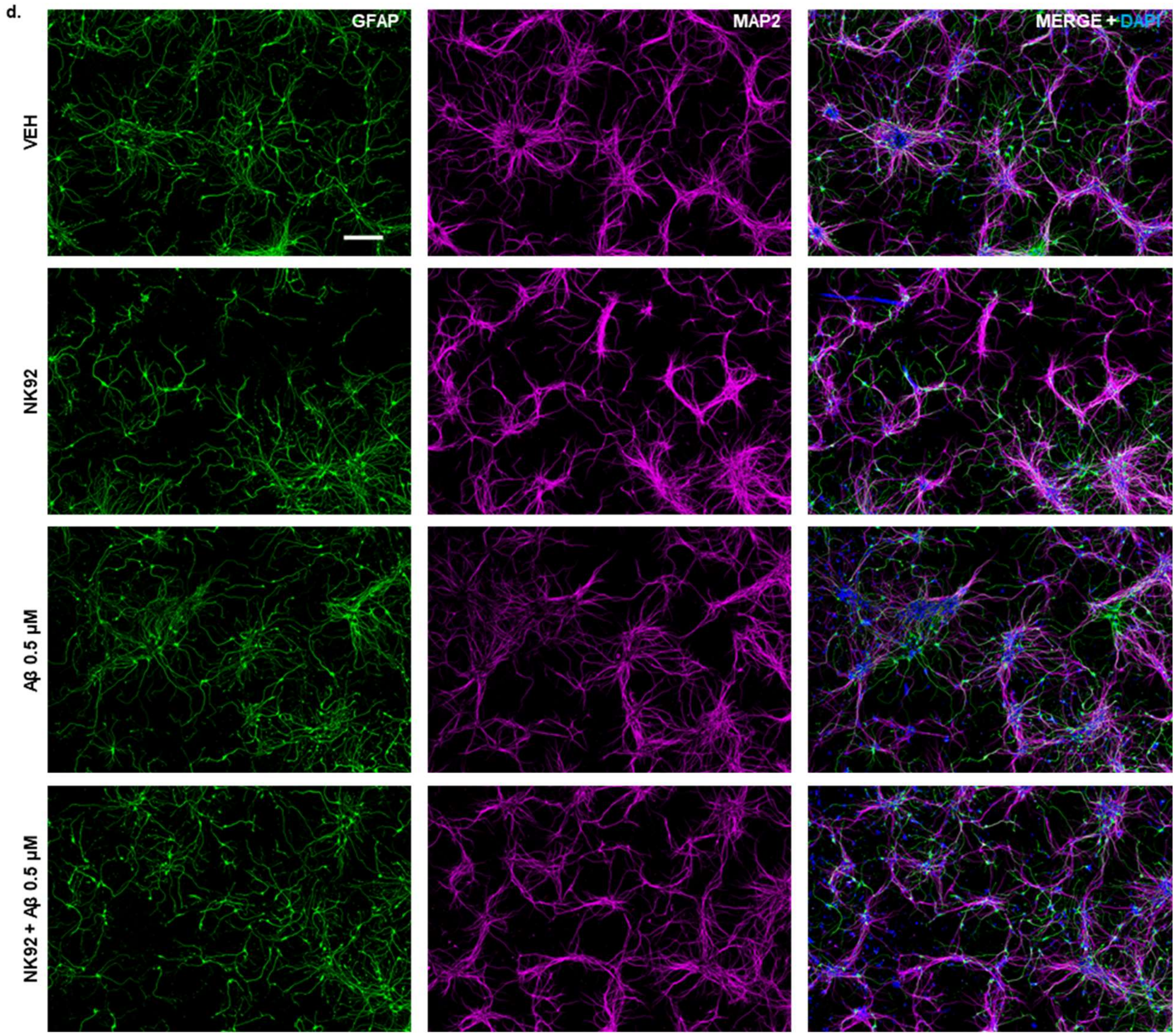


**Sup. Fig. 3d.** Tile scan images of 43 DIV human mixed primary neuron and astrocyte cultures for NK92 experiments. Only GFAP, MAP2, and DAPI channels are shown. GFAP and MAP2 identify astrocytic and neuronal processes, respectively. Merged images include DAPI to identify nuclei. 40x objective with 1x zoom. Scale bar = 200 µm.

**e.**

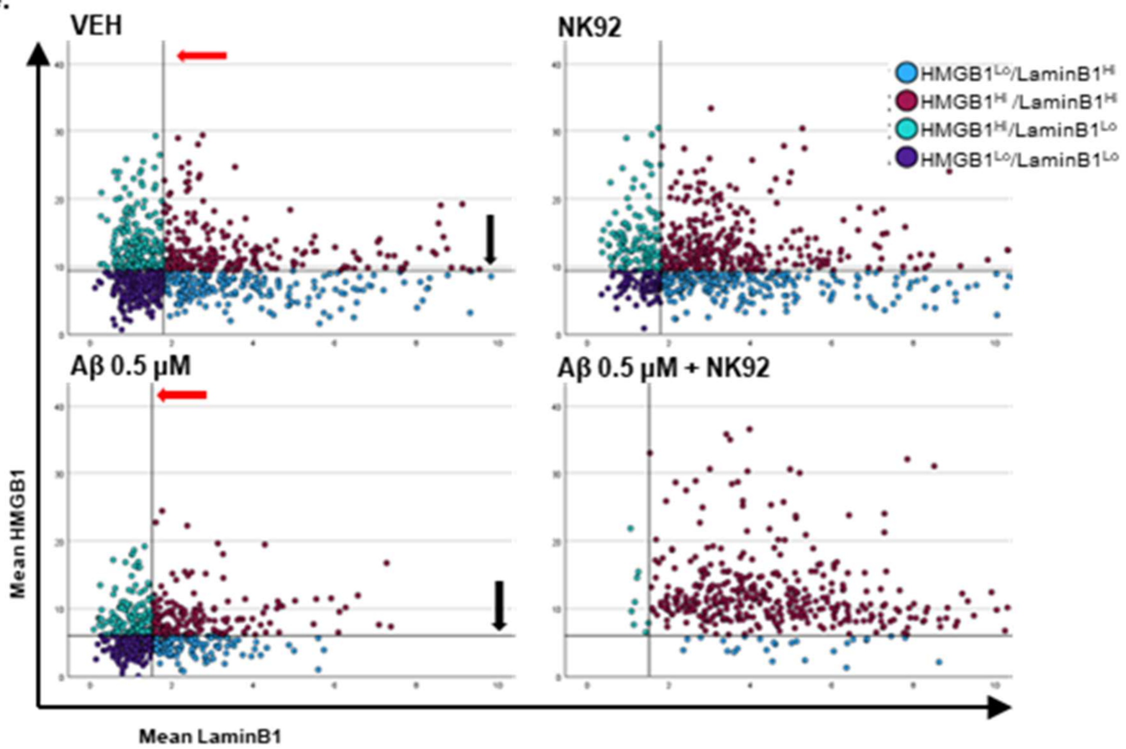


**Sup. Fig. 3e.** Scatter plot of the HMGB1 and LaminB1 mean signal for neurons of VEH, NK92, Aβ, and Aβ + NK92 treatment groups for one replicate (see Sup. Fig. 3f. for the other two replicates). First, $HMGB1^{Lo}/LaminB1^{Lo}$ populations are identified in VEH and Aβ-treated cells. To identify $HMGB1^{Lo}/LaminB1^{Lo}$ populations, cells are separated by the intersection of the medians of LaminB1 (red arrows) and HMGB1 (black arrows). The intersection of the medians results in 4 quadrants identifying the percentage of cells in $HMGB1^{Lo}/LaminB1^{Hi}$ (light blue), $HMGB1^{Hi}/LaminB1^{Hi}$ (red), $HMGB1^{Hi}/LaminB1^{Lo}$ (green), and $HMGB1^{Lo}/LaminB1^{Lo}$ (dark blue) quadrants. The medians of the VEH and Aβ groups are then used on NK92 and Aβ+NK92 groups, respectively, to generate quadrants based on their reference groups. The quadrants for $HMGB1^{Lo}/LaminB1^{Lo}$ populations contain putative senescent-like cells simultaneously expressing low LaminB1 and HMGB1 levels. The specific elimination of these senescent-like cells can be detected indirectly by assessing changes in the percentage of $HMGB1^{Lo}/LaminB1^{Lo}$ populations (dark blue) after NK92 co-culture. The fold increase in the percentage of VEH $HMGB1^{Lo}/LaminB1^{Lo}$ cells (dark blue) over the percentage of $HMGB1^{Lo}/LaminB1^{lo}$ cells (dark blue) in NK92 groups represents the loss of $HMGB1^{Lo}/LaminB1^{Lo}$ from VEH to NK92 (VEH→NK92). Likewise, the fold increase in the percentage of $HMGB1^{Lo}/LaminB1^{Lo}$ cells (dark blue) in Aβ compared to Aβ+NK92 groups (Aβ→Aβ+NK92) reflects the loss of populations potentially containing Aβ-treated senescent-like cells. The fold change of $HMGB1^{Lo}/LaminB1^{Lo}$ cells in VEH→NK92 and Aβ→Aβ+NK92 are then average normalized replicate by replicate and statistically compared.

f.

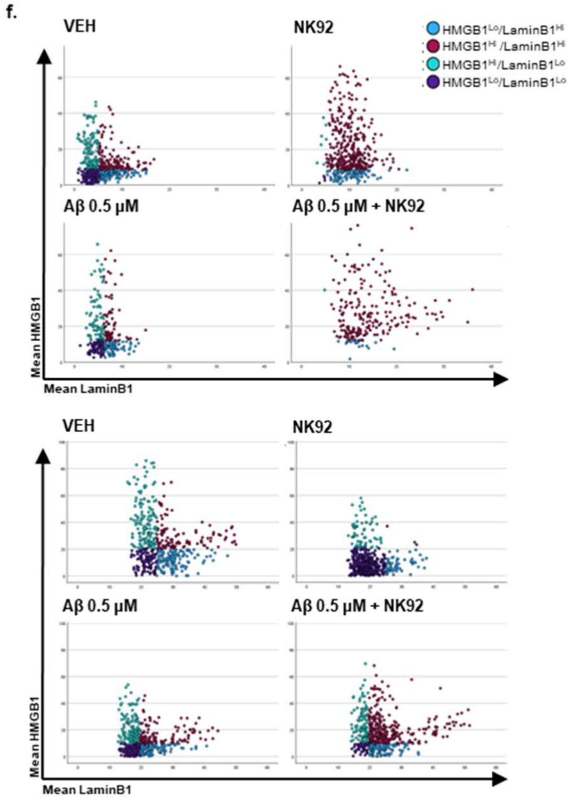


**Sup. Fig. 3f.** Scatter plots of HMGB1$^{Lo}$/LaminB1$^{Lo}$ neuron populations (dark blue) for VEH, NK92, Aβ, and Aβ + NK92 for remaining replicates not shown in Sup. Fig. 3e, obtained as described in Sup. Fig. 3e.

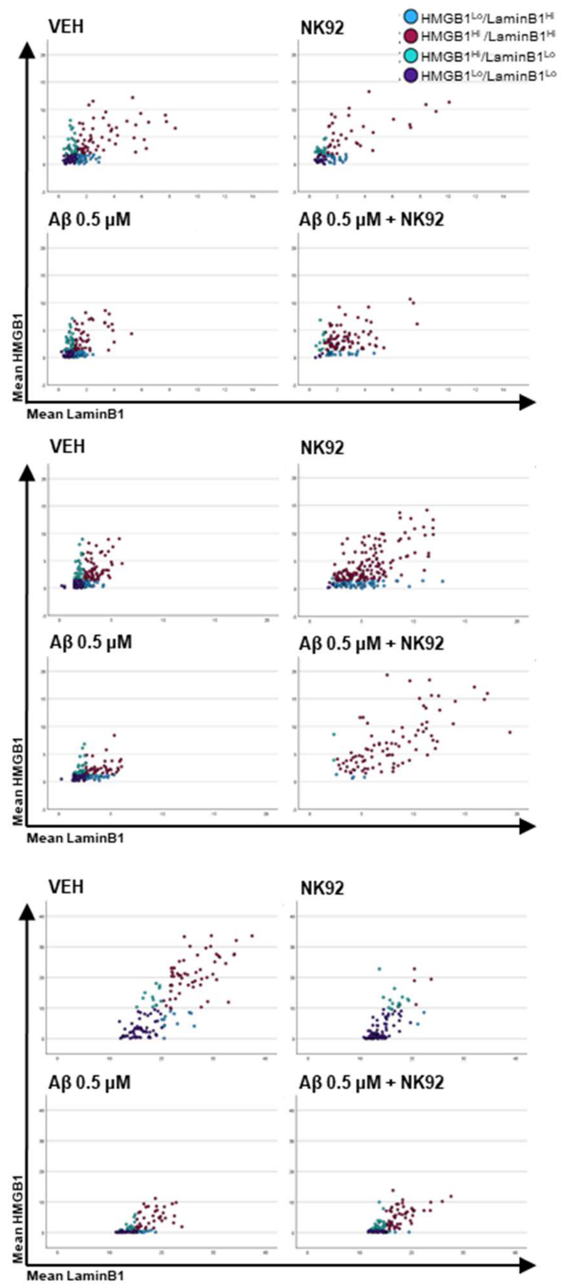


**Sup. Fig. 4.** Scatter plots of $HMGB1^{Lo}/LaminB1^{Lo}$ astrocyte populations (dark blue) for VEH, NK92, Aβ, and Aβ + NK92 conditions for all replicates, obtained obtained as described in Sup. Fig. 3e.

.

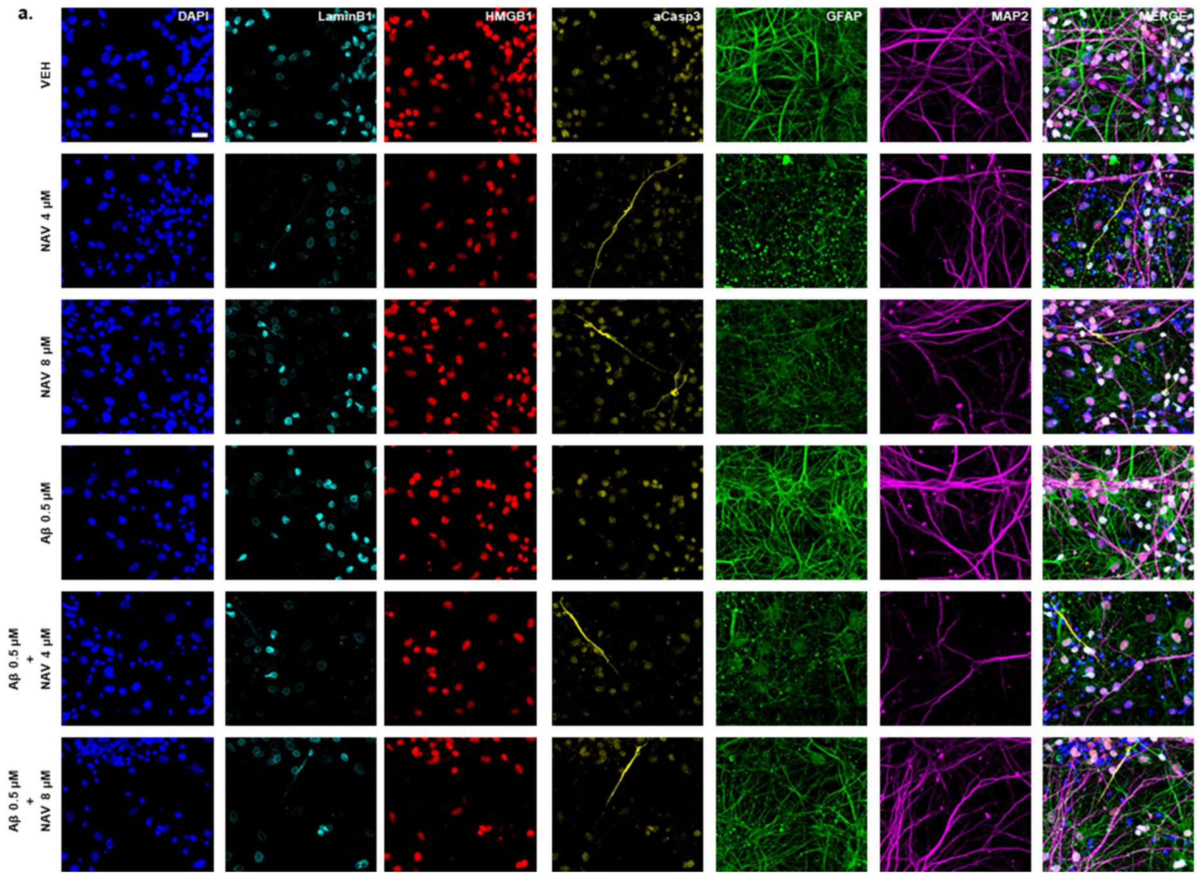


**Sup. Fig. 5a.** Representative linearly unmixed confocal spectral scans of 43 DIV mixed primary human and astrocyte cultures from NAV 4 and 8 µM treatment experiments, stained for DAPI, LaminB1, HMGB1, aCasp3, GFAP, and MAP2. Treatments with NAV reveal MAP2/GFAP-negative aCasp3-positive cell remnants exhibiting elongated processes resembling neuronal processes. NAV treatment conditions also show thinning of MAP2 dendritic arborization. A reduction in GFAP-positive processes is also observed, accompanied by the presence of prominent GFAP-positive puncta, which may represent remnants of astrocytic processes. 40x objective with 1x zoom. Scale bar = 20 µm.

**b.**

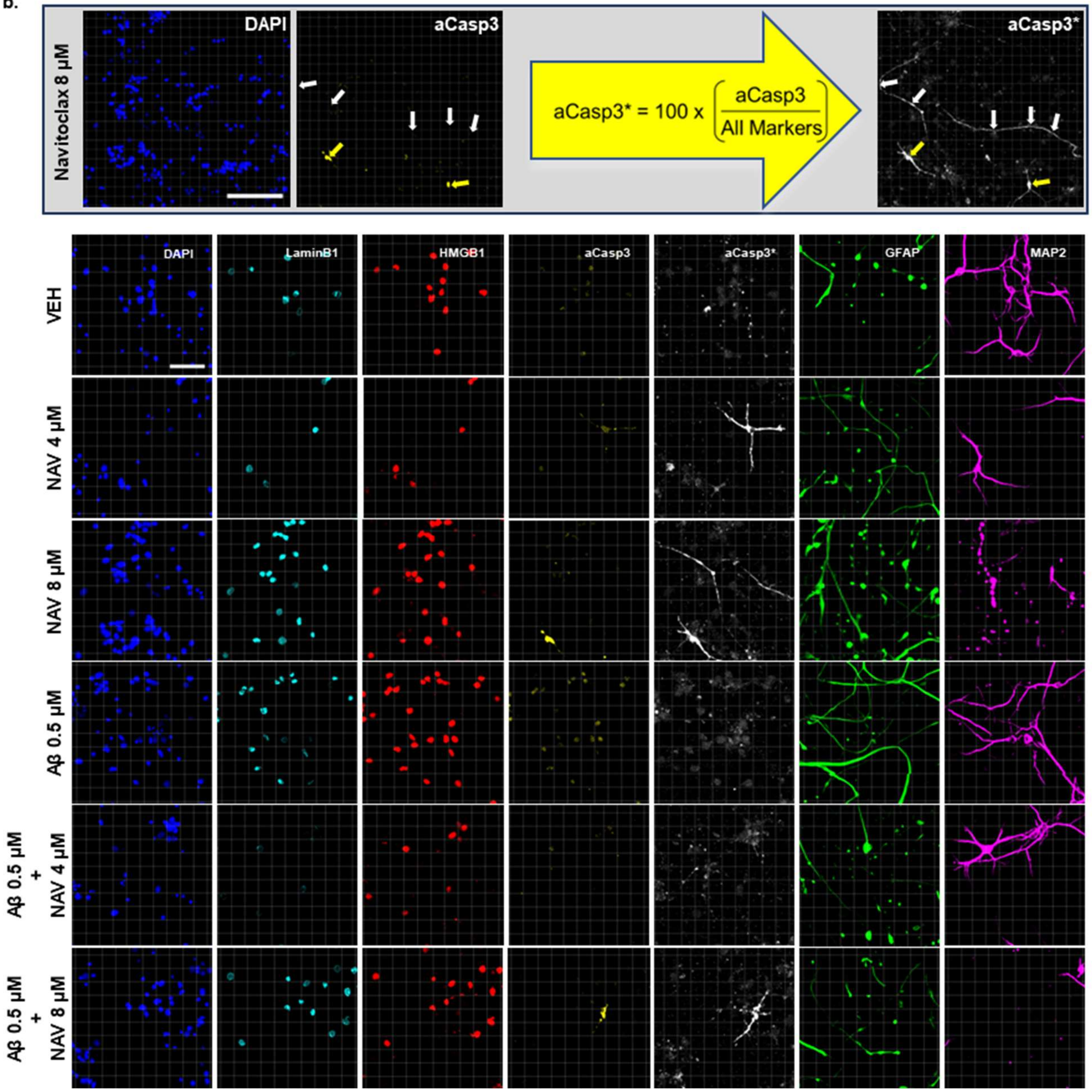


**Sup. Fig. 5b. Grey box.** Images of NAV 8 μM treatments illustrating the channel arithmetic operation (yellow arrow) to process aCasp3 into aCasp3*, which adjusts aCasp3 signal in inverse proportion to the remaining channels. This adjustment amplifies the dim aCasp3 signal that is negative for the rest of the markers, uncovering axon-like processes (indicated by white arrows) and enhancing the visualization of apoptotic cell remnants (indicated by yellow arrows). Further details are provided in Materials and Methods. 40x objective with 1x zoom. Scale bar = 50 μm. **Bottom rows.** Representative confocal images of all treatment conditions with side-by-side comparison of aCasp3 and aCasp3* channels. 40x objective with 1x zoom. Scale bar = 25 μm.

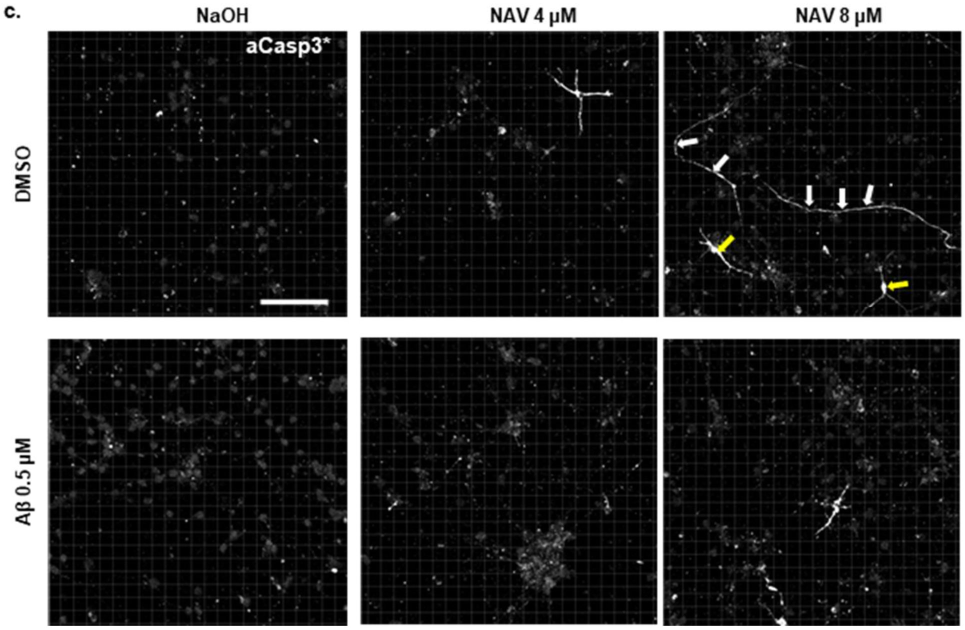


**Sup. Fig. 5c.** Representative confocal images of aCasp3* for all treatment conditions. aCasp3* is processed from aCasp3 as explained in Sup. Fig. 5b. In NAV 8 µM treatment images, white arrows identify potential axonal aCasp3 remnants and yellow arrows identify cell body remnants. 40x objective with 1x zoom. Scale bar = 50 µm.

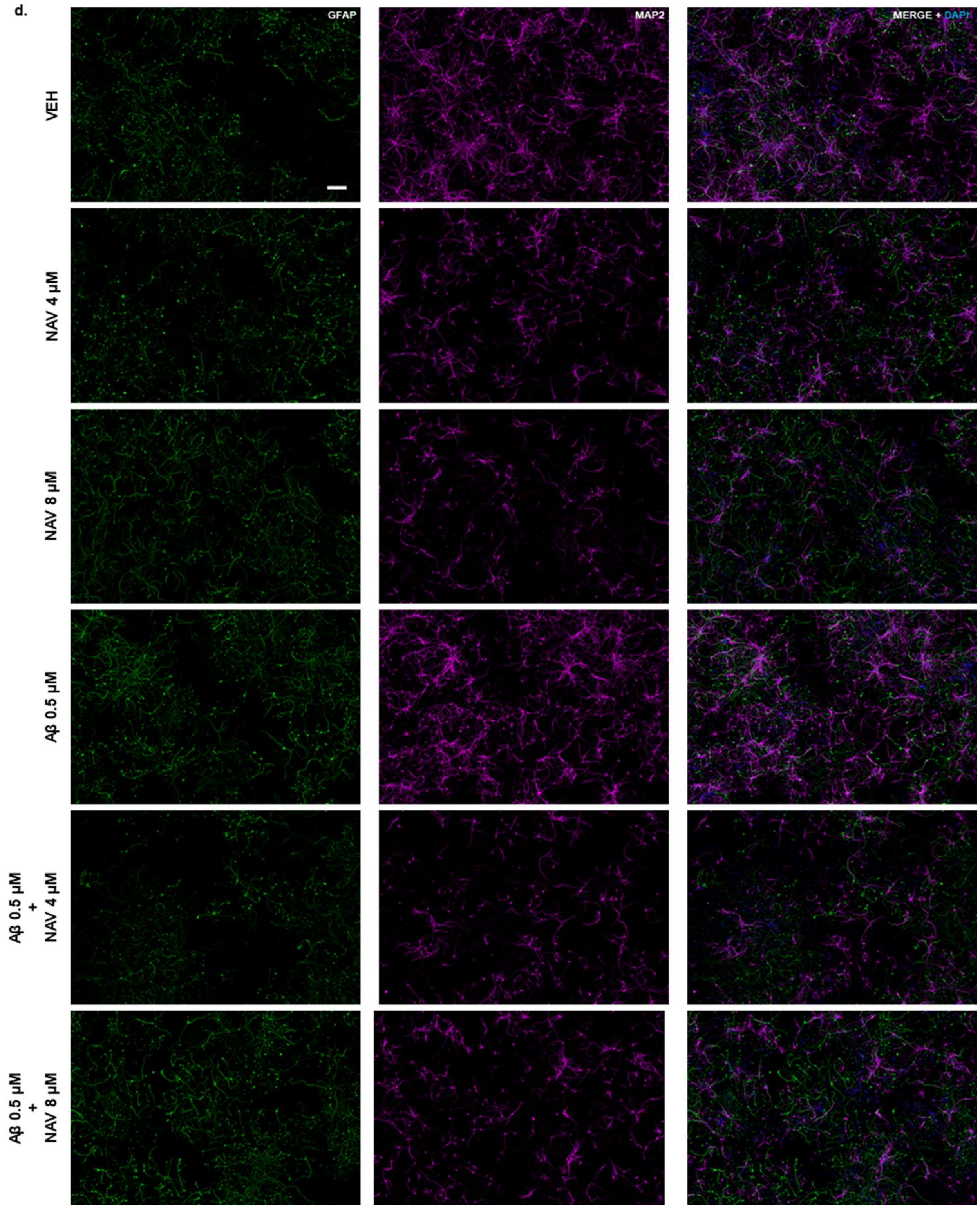


**Sup. Fig. 5d.** Representative tile-scan images of 43 DIV human mixed primary neuron and astrocyte cultures for NAV 4 & 8 µM experiments. GFAP and MAP2 identify astrocytic and neuronal processes, respectively. Merged images include DAPI to identify nuclei. NAV treatments at 4 and 8 µM present reduced MAP2 positive arborization irrespective of Aβ treatment. 40x objective with 1x zoom. Scale bar = 200 µm.

**e.**

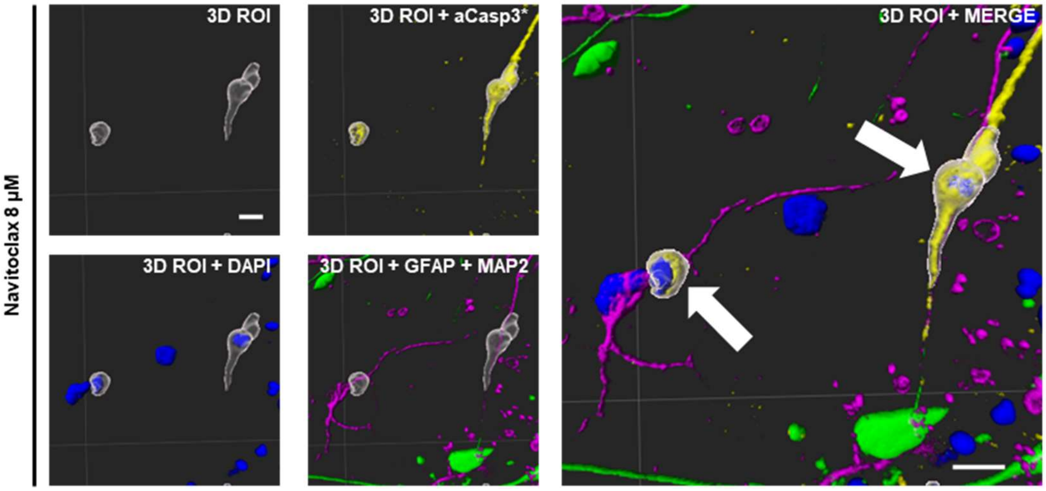


**Sup. Fig. 5e.** Representative 3D regions of interest (3D ROI) of apoptotic cells obtained from confocal images of NAV 8 µM treatment. 3D ROI image (small image top, left) shows examples of two apoptotic cell remnant ROIs without markers. 3D ROI + aCasp3* (small image top, right) and 3D ROI + DAPI (small image bottom, left) show colocalization of 3D ROIs with aCasp3* and DAPI staining, respectively. 3D ROI + GFAP + MAP2 image (small images bottom, right) shows lack of colocalization with GFAP and MAP2 immunostaining. 3D ROI + MERGE image (large image) shows apoptotic cell remnant 3D ROIs (white arrows) with aCasp3*, DAPI, GFAP, and MAP2 channels merged. To obtain apoptotic cell 3D ROIs, the aCasp3* channel described in Sup. Fig. 5b was used to generate 3D ROI on aCasp3*-positive apoptotic cell remnants that were negative for MAP2 and GFAP (data not shown). The DAPI channel was used to generate 3D ROI on nuclei displaying pyknosis or karyorrhexis, characteristic of cell death (data not shown). The overlapping 3D ROI generated from aCasp3* and DAPI were used to create new 3D ROI of apoptotic cells shown as exemplified in the images. Additional details in Materials and Methods. 40x objective with 1x zoom. Scale bar = 10 µm.

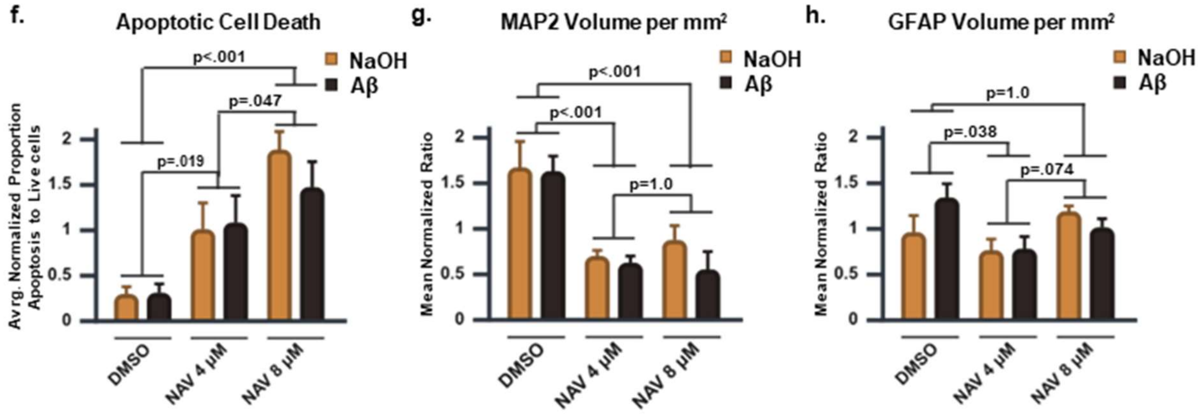


**Sup. Fig. 5. Preliminary assessment of NAV 4 & 8 µM on mixed human primary neurons and astrocytes in Aβ model of AD. (f)** Bar graph of apoptotic cell death showing non-selective elimination by NAV **(g)** Bar graph of MAP2 per mm$^2$ showing a non-selective reduction of the volume of neuronal processes. **(h)** Bar graph of GFAP per mm$^2$ showing a reduction of astrocytic processes at low but not high NAV concentrations. Bonferroni-adjusted all pairwise comparisons are carried out on the marginal means for NAV treatments. Bar charts display all group combinations instead of the marginal means to visualize trends. Significance, $p < .05$. 3x Replicates. Graphs represent mean ± s.e.m.

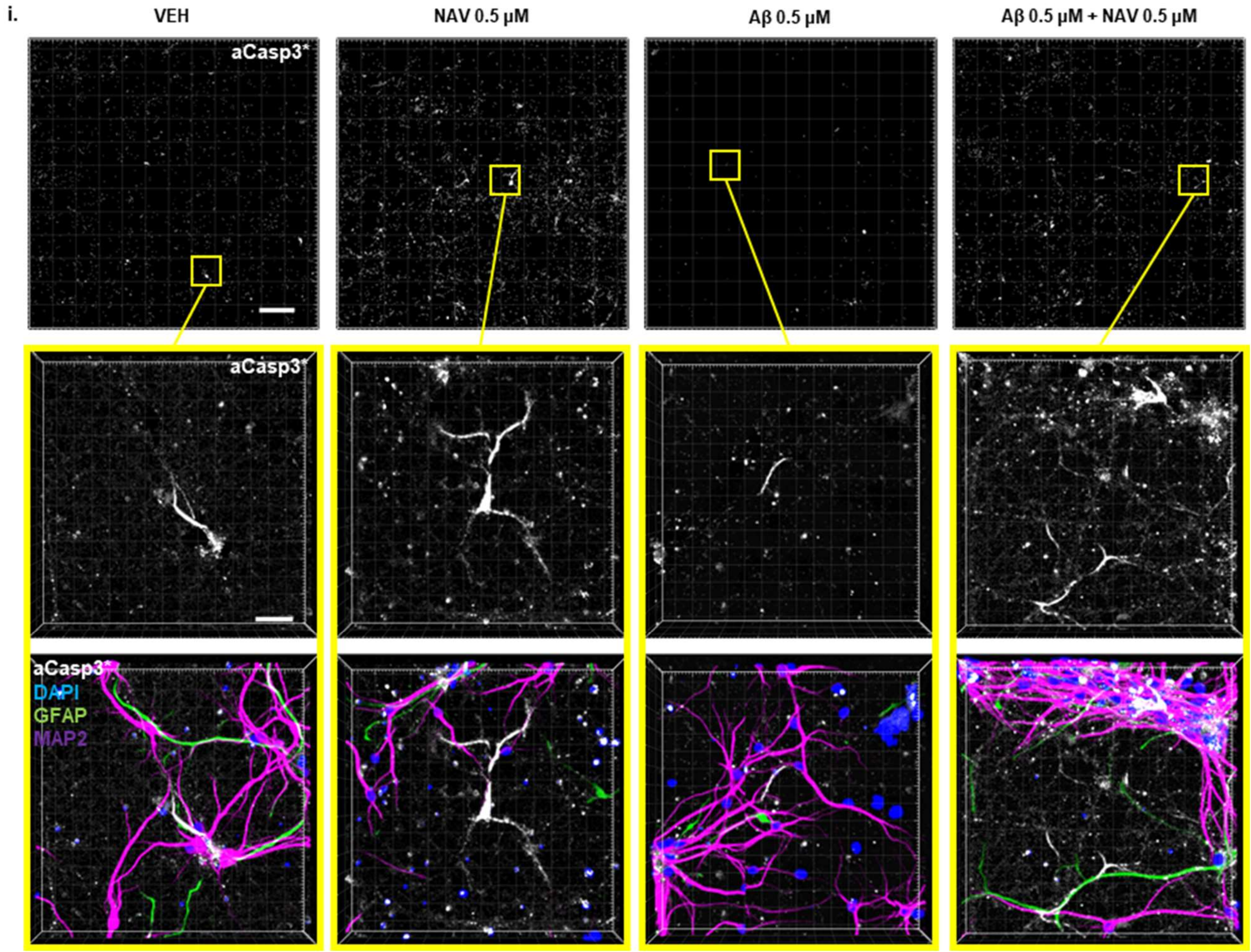


**Sup. Fig. 5i.** Low NAV (0.5 μM) concentration representative 3D confocal tile scan images of aCasp3* (top row) with detail of aCasp3* (center row) and additional DAPI, GFAP, and MAP2 merged image (bottom row) for all treatment conditions. aCasp3* is processed from aCasp3 as explained in Sup. Fig. 5b. Tile-scan images (top row) show reduced aCasp3* signal in VEH and Aβ treatments that is increased in NAV treatments. Magnified images of aCasp3* (center row) of tile-scans (top row) display morphological traits of aCasp3* apoptotic cell remnants. Bottom row images show lack of colocalization of aCasp3* with DAPI, GFAP, and MAP2. 40x objective with 1x zoom. Tile scans (top row) scale bar = 300 μm. Detail images (center and bottom rows) scale bar = 40 μm j.

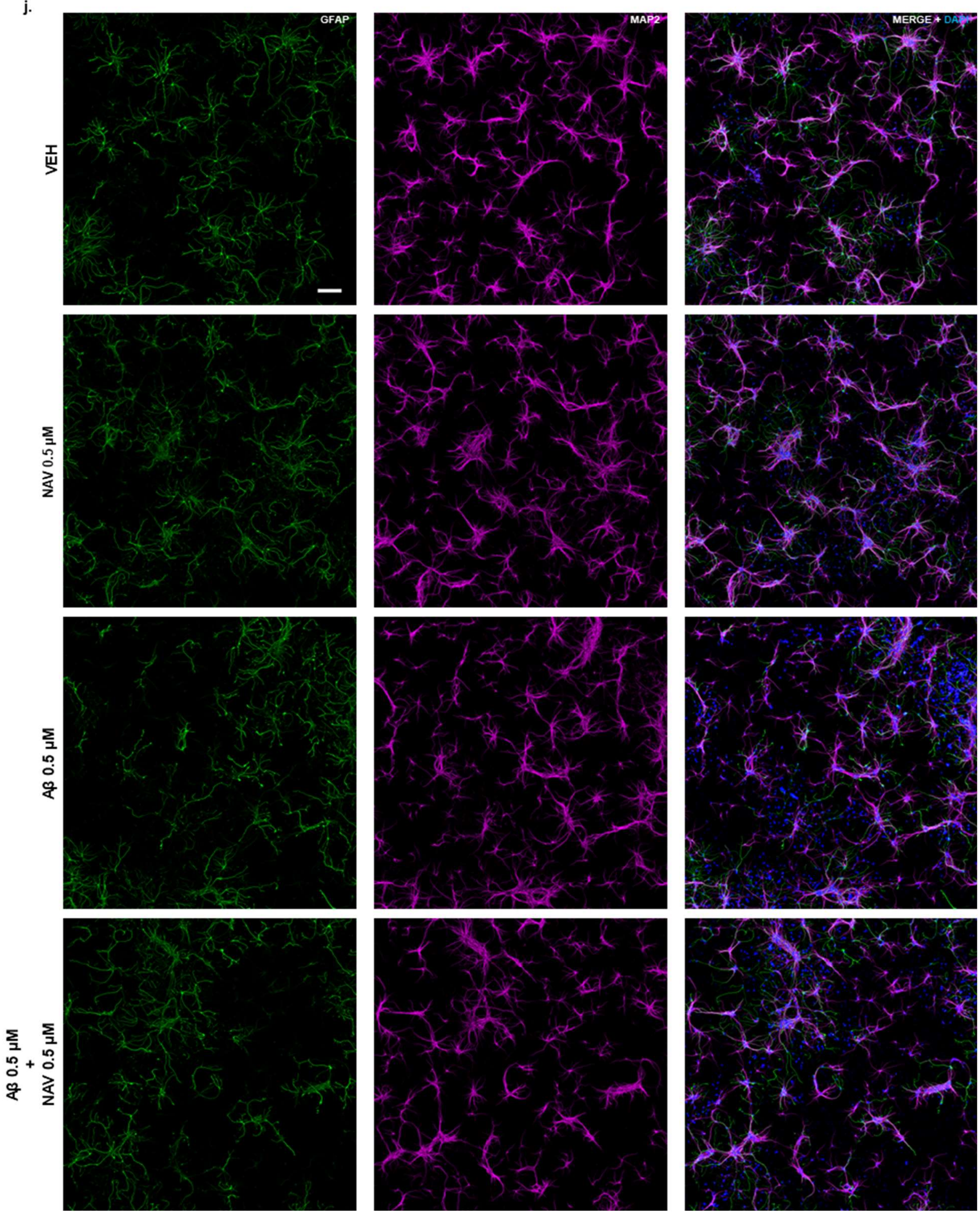


**Sup. Fig. 5j.** Representative tile scan images of 43 DIV mixed human primary neuron and astrocyte cultures for NAV 0.5 µM experiments. GFAP and MAP2 identify astrocytic and neuronal processes, respectively. Merged images include DAPI to identify nuclei. Unlike in NAV treatments at 4 and 8 µM (Sup. Fig. 5d), there are no readily evident differences in MAP2 or GFAP-positive processes. 40x objective with 1x zoom. Scale bar = 200 µm.

**k.**

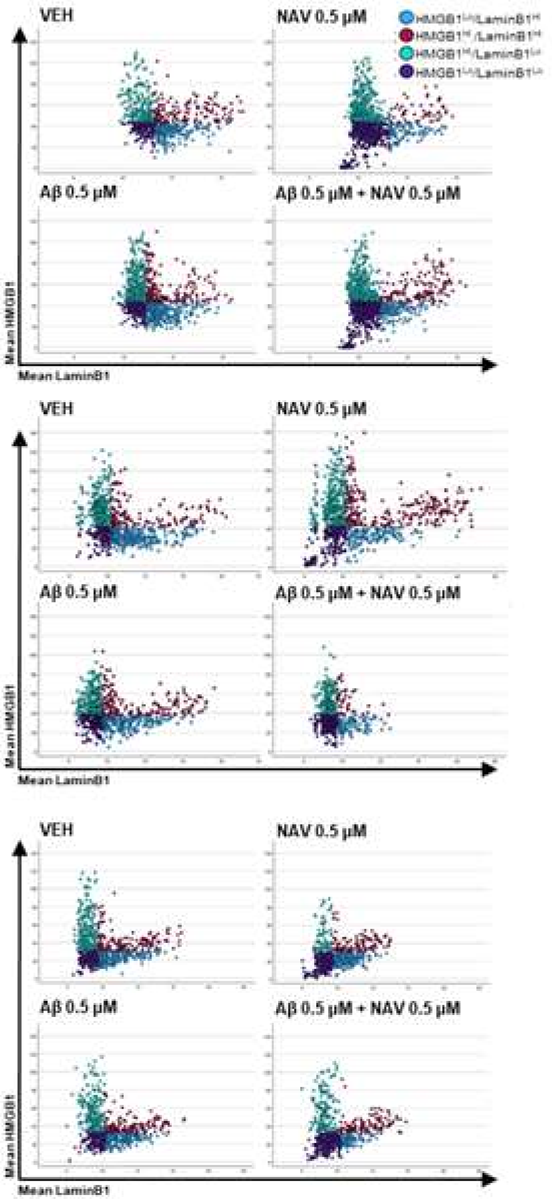


**Sup. Fig. 5k.** Scatter plots of HMGB1$^{Lo}$/LaminB1$^{Lo}$ neuron populations (dark blue) for VEH, NAV, Aβ, and Aβ + NAV conditions for all replicates, obtained as described in Sup. Fig. 3e.

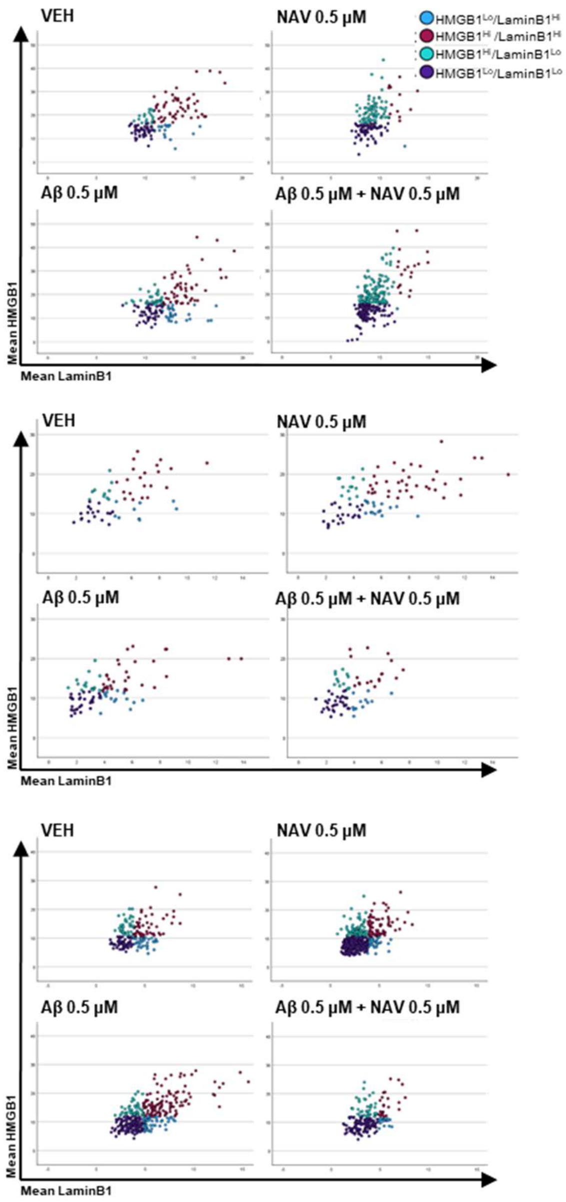


**Sup. Fig. 6.** Scatter plots of HMGB1$^{Lo}$/LaminB1$^{Lo}$ astrocyte populations (dark blue) for VEH, NAV, Aβ, and Aβ + NAV conditions for all replicates, obtained as described in Sup. Fig. 3e.

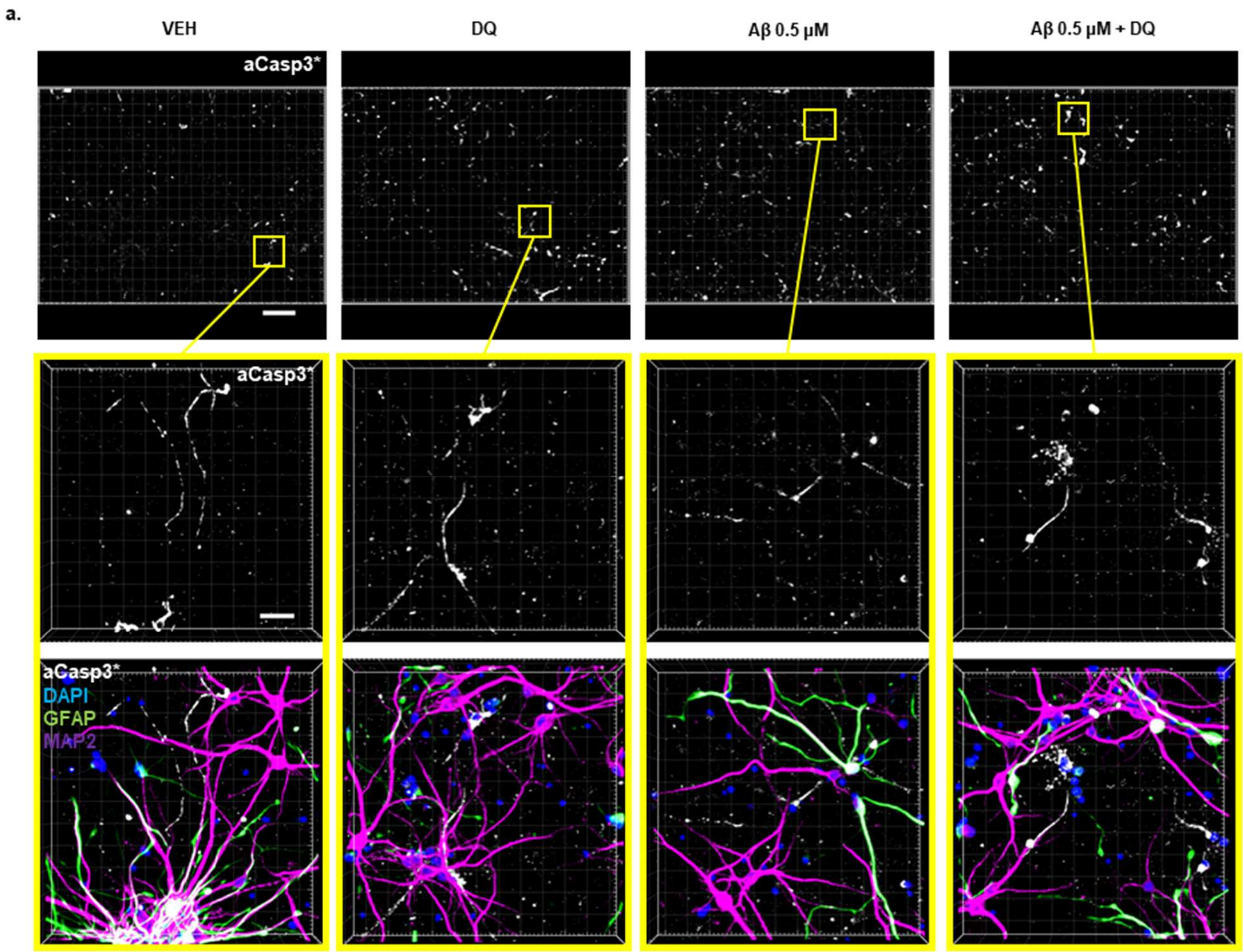


**Sup. Fig. 7a.** Representative confocal tile scan images of aCasp3* (top row) with detail of aCasp3* (center row) and additional DAPI, GFAP, and MAP2 merged image (bottom row) for all treatment conditions. aCasp3* is processed aCasp3 as explained in Sup. Fig. 5b. Tile scan images (top row) and details (center row) of aCasp3* do not show evident differences in aCasp3* across treatment conditions. 40x objective with 1x zoom. Tile scans (top row) scale bar = 300 μm. Detail images (center and bottom rows) scale bar = 30 μm

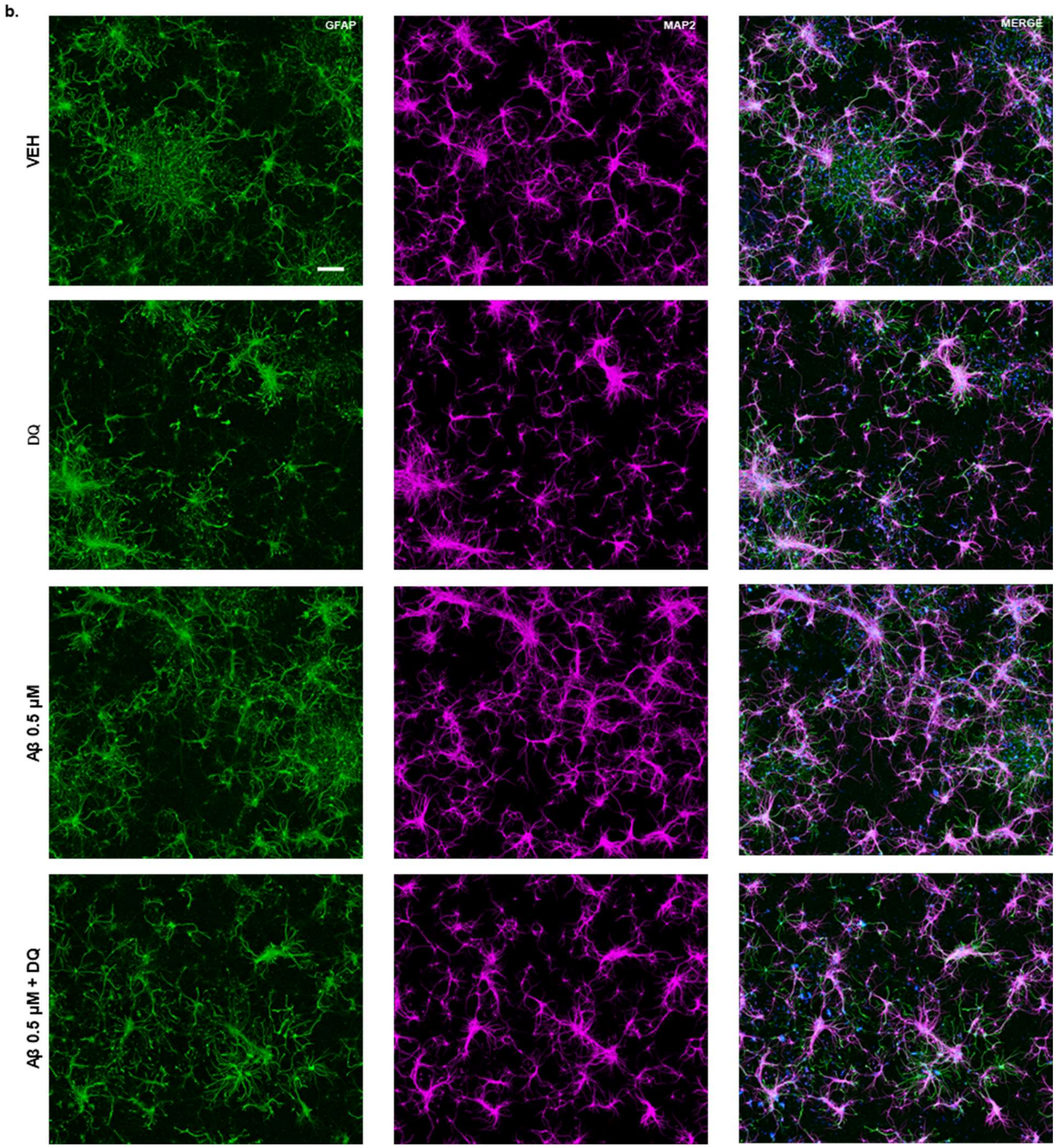


**Sup. Fig. 7b.** Representative tile scan images of 43 DIV mixed human primary neuron and astrocyte cultures for DQ experiments. GFAP and MAP2 identify astrocytic and neuronal processes, respectively. Merged images include DAPI to identify nuclei. There are no readily evident differences in MAP2 or GFAP processes between treatment conditions. 40x objective with 1x zoom. Scale bar = 200 µm.

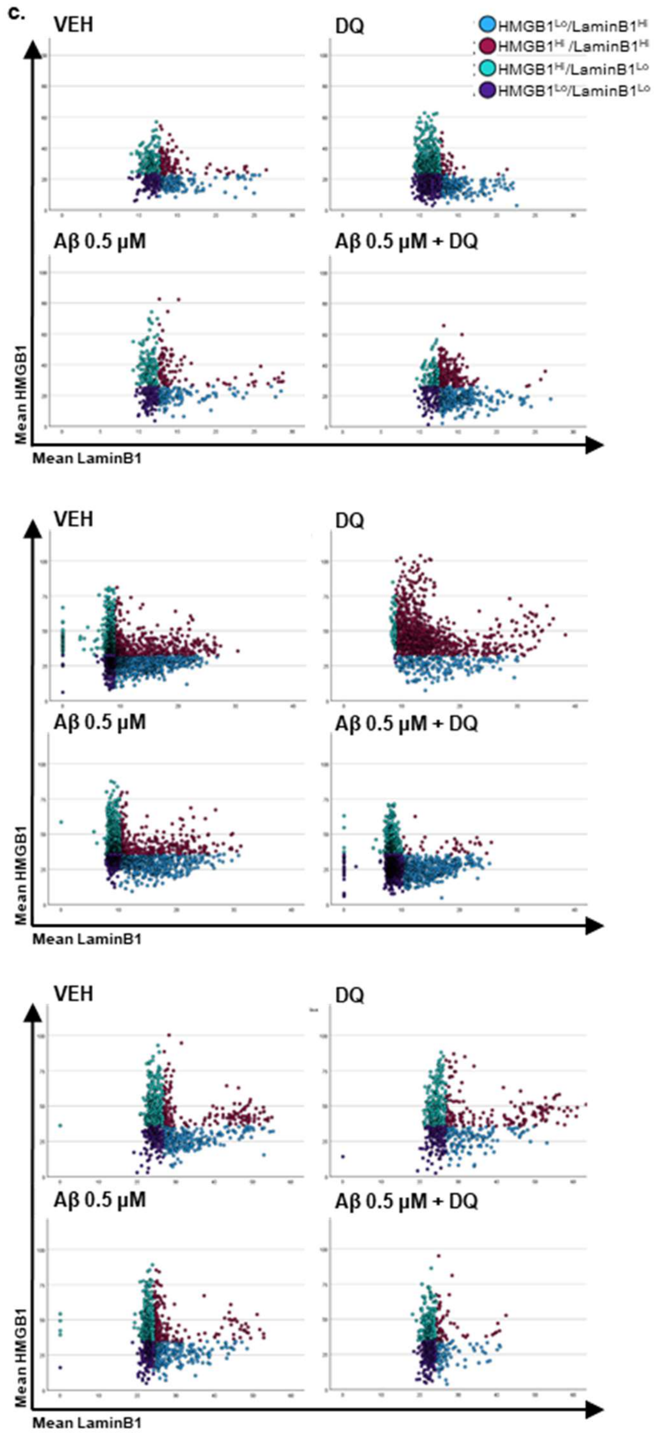


**Sup. Fig. 7c.** Scatter plots of HMGB1$^{Lo}$/LaminB1$^{Lo}$ neuron populations (dark blue) for VEH, DQ, Aβ, and Aβ + DQ conditions for all replicates, obtained as described in Sup. Fig. 3e.

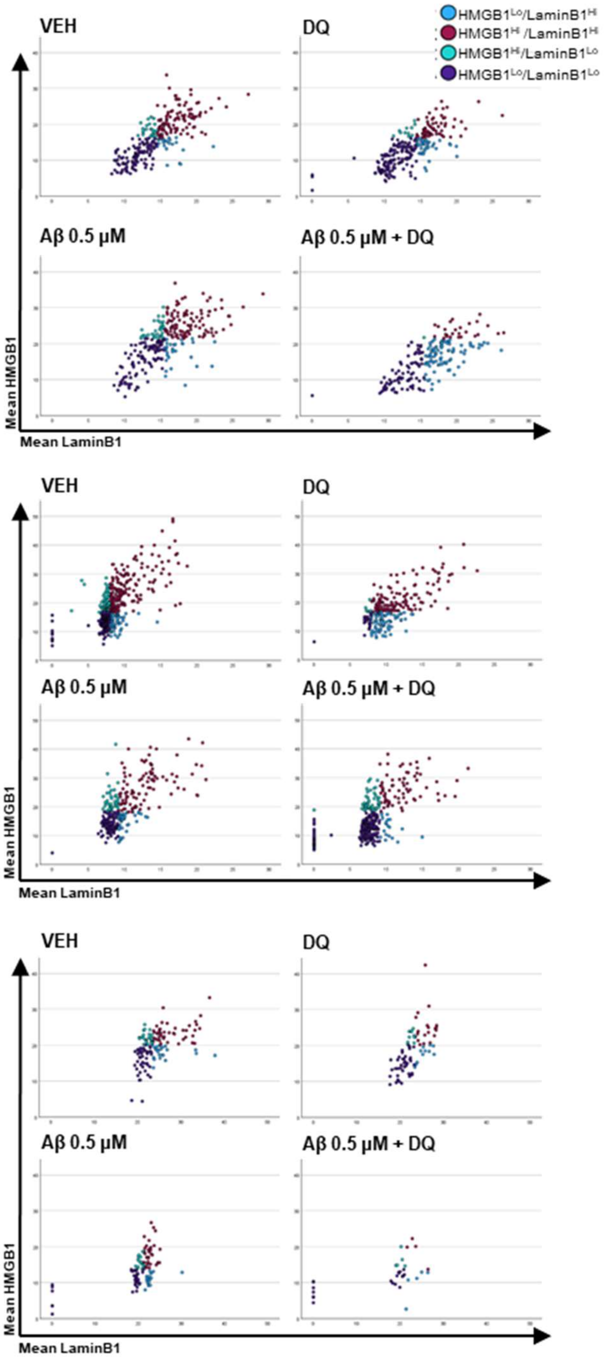


**Sup. Fig. 8.** Scatter plots of HMGB1$^{Lo}$/LaminB1$^{Lo}$ astrocyte populations (dark blue) for VEH, DQ, Aβ, and Aβ + DQ conditions for all replicates, obtained as described in Sup. Fig. 3e.
